# Myeloid reprogramming by JAK inhibition enhances checkpoint blockade therapy

**DOI:** 10.1101/2022.06.24.497435

**Authors:** Jaroslav Zak, Isaraphorn Pratumchai, Brett S. Marro, Kristi L. Marquardt, Reza Beheshti Zavareh, Luke L. Lairson, Michael B. A. Oldstone, Judith A. Varner, Veronika Bachanova, John R. Teijaro

**Author notes:** Janssen Research and Development, San Diego, 92121 CA. Trotana Therapeutics, 5555 Oberlin Drive, Suite 100, San Diego, CA. equal contribution (Jaroslav Zak, Isaraphorn Pratumchai, Brett S. Marro).

## Abstract

Unleashing anti-tumor T cell activity by checkpoint inhibition is effective in many cancer patients but clinical response rates remain limited. Myeloid derived suppressor cells erode antitumor lymphocyte numbers and function, and correlate with resistance to checkpoint inhibitors. By screening small molecule libraries, we identified JAK inhibitors’ ability to rescue T cell function. Despite its documented immune suppressive properties, the prototypical JAK inhibitor ruxolitinib enhanced the efficacy of immune checkpoint blockade in cancer. This effect correlated with loss of suppressive gene expression, and acquisition of immunostimulatory molecular markers and T cell stimulatory activity in myeloid cells. In preclinical models, ruxolitinib significantly improved the function and increased the total numbers of activated tumor-infiltrating NK and CD4 T cells compared to checkpoint blockade alone and the efficacy was conditional on granulocytic cells. In addition to myeloid reprogramming in the tumor, ruxolitinib blunts G-CSF signaling in the bone marrow to prevent expression of suppressive and chemotaxis genes in neutrophils. In a clinical trial of Hodgkin lymphoma patients resistant to checkpoint inhibitors, treatment with ruxolitinib significantly reduced neutrophil-to-lymphocyte ratios and levels of suppressive markers in myeloid cells but increased numbers of cytokine-producing T cells. These results support the therapeutic potential of JAK inhibition in combination with checkpoint inhibitors in cancer and highlight the potential of reshaped myeloid immunity to improve immunotherapy.

One sentence summary: Ruxolitinib reshapes myeloid immunity to synergize with checkpoint inhibitors

## Introduction

Immune control of cancer and response to immunotherapy are hampered by diverse, partially redundant, adaptive immunosuppressive effects mediated by both cancer and non-cancer cells. Suppressive myeloid cells are present in many tumor types, associated with lymphocyte dysfunction and a poor response to immune checkpoint inhibitors (*1, 2*). The pan immune suppressive state generated during persistent infection closely resembles that observed in cancer (*3, 4*). In fact, the important cellular and molecular features of immune suppression including T cell exhaustion and suppressive myeloid cells are shared between cancer and persistent viral infection, leading to fundamental discoveries such as T cell reactivation by PD1 blockade in the lymphocytic choriomeningitis virus (LCMV) Clone 13 model (*5-7*).

Targeting of myeloid derived suppressor cells (MDSCs) and tumor associated macrophages enhances response to checkpoint inhibitors in cancer models. Interestingly, emerging evidence suggests that suppressive activity may be associated with stimulus-dependent states in all myeloid lineages (*8, 9*). The upstream stimuli that drive suppressive programming of myeloid cells offer a promising therapeutic opportunity. Multiple signal transducer of activators of transcription (STAT) transcription factors and the JAK1/2-dependent cytokines G-CSF, GM-CSF and IL-6 are implicated in suppressive programming of MDSCs (*9, 10*). Therapeutics specifically targeting these cells have yet to gain clinical approval (*11*) and conflicting effects of Janus kinase (JAK) inhibitors on MDSCs have been reported (*12-14*). Therefore, clinically applicable approaches to targeting MDSCs for cancer therapy are highly sought.

The JAK/STAT pathway plays a central role in activating transcriptional programs in responses to dozens of soluble mediators including cytokines, growth factors and interferons (*15*). As such, JAK/STAT signaling plays integral roles in cell growth, differentiation and survival. Several small molecule JAK inhibitors have been approved in the clinic and are predominantly used to treat myeloproliferative and autoimmune diseases (*16*). Despite known genetic links between JAK mutations and cancer, mainstream use of JAK inhibitors for cancer treatment has been impeded by JAK inhibitors’ immune suppressive properties. Here we demonstrate that rather than suppress essential anti-tumor immunity, small molecule JAK inhibition synergizes with checkpoint therapy to reshape the myeloid cell compartment and enhance NK and T cell responses. Importantly, conjunctive JAK inhibition and checkpoint blockade enhanced tumor control in both mice and humans.

## Results

### JAK inhibitors rescue function of exhausted T cells

We developed a screening strategy to discover small molecules that rescue T cell exhaustion (*17*). The assay relies on flow-cytometric detection of YFP as a reporter of IFN-γ expression within hypofunctional CD8 or CD4 T cells isolated from mice persistently infected with LCMV Clone 13 (Cl13). Persistent infection with Cl13 is characterized by a broadly immunosuppressive environment featuring T cell exhaustion and suppressive activity by myeloid cells (*6*). Having screened the ReFrame collection of ∼12,000 biologically active compounds using the assay, we sought to determine if specific biomolecular targets were overrepresented in the set of hit compounds. We examined annotated compound-gene interaction curated by the Drug-Gene Interaction Database (*18*). The most significantly enriched target among compounds increasing both percentage and number of YFP^+^ CD8 T cells was GSK-3β, a known target whose inhibition reduced the expression of inhibitory receptors on CD8 T cells and improved antitumor responses (**Figure 1A-B**) (*19*). We found an enrichment of JAK inhibitors (JAKi) among hit compounds (**Figure 1B-C**). Enrichment of JAKi was statistically significant among hits that increased the number and percentage of YFP^+^ CD8 T cells, and among hits that increased either parameter alone (**Figure 1C**). These inhibitor compounds span multiple chemotypes, arguing against a common off-target (**Figure S1A**). The top 5 JAKi by z-score (BMS-911543, ruxolitinib, decernotinib, R-333 and AZD-1480) were re-obtained and their activity re-tested at a range of concentrations (**Figure S1B**). Although all compounds showed some activity in the validation assay, only BMS911543 and ruxolitinib could match or surpass the effect of aPDL1 (**Figure S1B**). To rule out YFP-dependent confounding effects we further validated the activity of ruxolitinib by assessing IFN-γ protein using intracellular staining in place of the YFP reporter (**Figure 1D**). To determine which kinase targets were specifically important for compound activity, a panel of 130 commercially available kinase inhibitors was assayed using the YFP-reporter high throughput platform. Many kinase inhibitors showed activity and the top hits were structurally diverse (**Figure 1E, S1C**). Interestingly, compounds inhibiting JAK1 and/or JAK2 were among the most potent in increasing the frequency of YFP^+^ CD8 T cells (**Figure 1F**).

**Figure 1.**
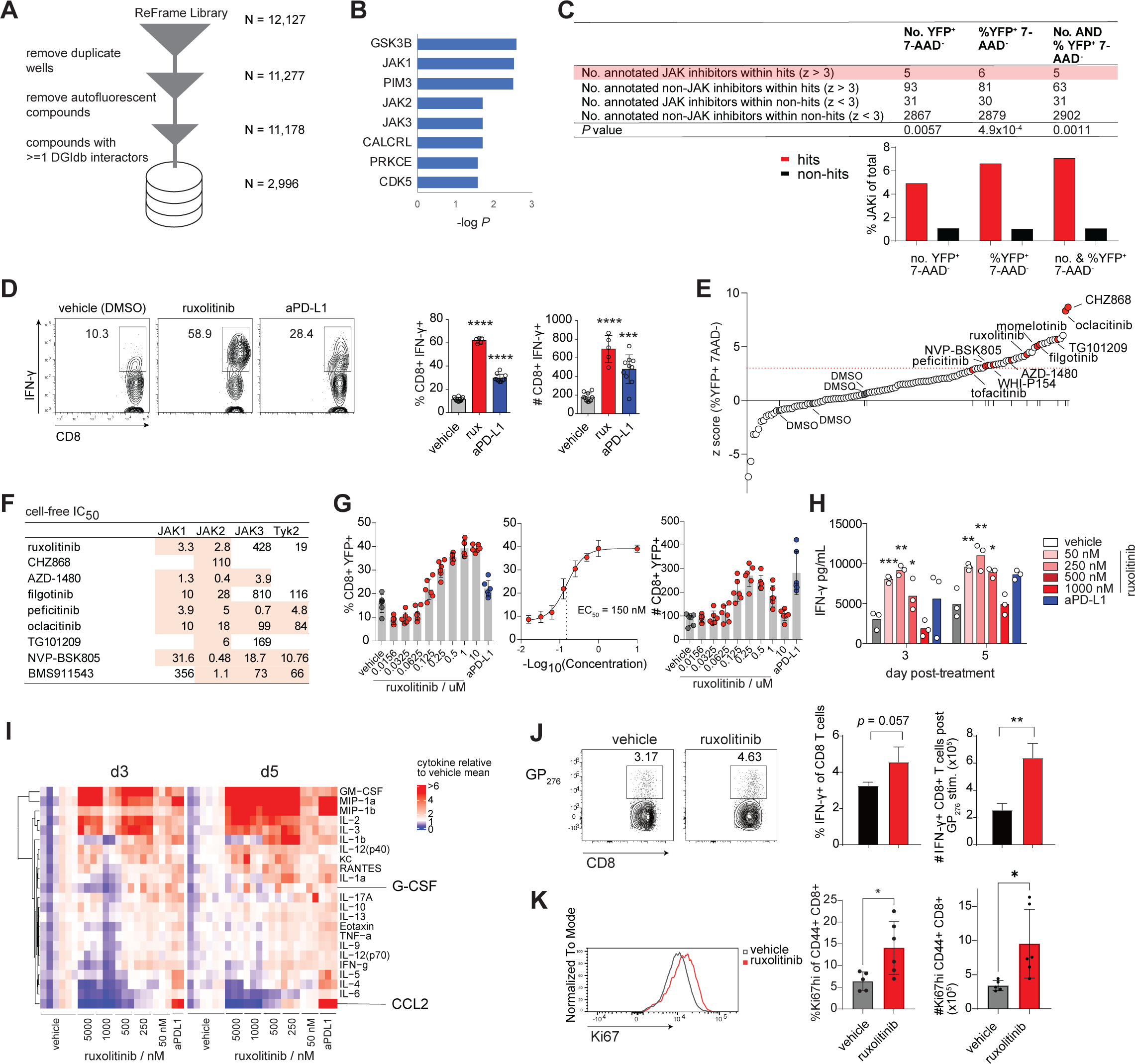
JAK1/2 inhibitors rescue proliferation of cytokine-producing CD8 T cells without impairing viral control. (A-C, E-H) Splenocytes from Cl13 infected IFNγ-IRES-YFP mice were harvested at 15 days post infection and cultured in the presence of LCMV peptides for 5 days (more details see Materials and Methods). (A) Preprocessing of ReFrame metadata for drug-target analysis, counts in right column indicate number of compounds at each step; (B) protein targets most highly enriched in hit vs non-hit compounds, targets were ranked by Fisher test p value and known drug metabolism enzymes were filtered out; (C) Summary of JAK inhibitor enrichment in hit vs non-hit groups using database-annotated compounds as described in panel A; (D) validation assay: effect of ruxolitinib and aPD-L1 treatment on endogenous IFN-γ labeled by intracellular staining, splenocyte culture conditions equivalent to primary assay with B6 mice substituted for YFP-IFN-γ mice; (E) ranking of compounds from kinase inhibitor library by z-score of %YFP^+^ 7-AAD^-^, JAKi marked in red, vehicle (DMSO only) controls highlighted in text; (F) cell-free IC_50_ values from JAK family inhibition assays of top JAKi hits from kinase inhibitor library, for list of references see Materials and methods; (G) dose response and EC50 calculation of ruxolitinib in enhancing %YFP^+^ of CD8 T cells in 5-day culture assay; (H) extracellular IFN-γ levels in supernatants of cultures with ruxolitinib or aPD-L1 or vehicle, Bio-plex immunoassay was used to measure IFN-γ; (I) extracellular cytokine levels 3 and 5 days post stimulation in splenocyte culture in the presence of varying concentrations of ruxolitinib, aPDL1 or vehicle, heatmap shows ratio of cytokine level to vehicle mean; (J-K) B6 mice were infected with Cl13 and treated with vehicle or ruxolitinib by daily gavage: (J) cytokine production in splenic CD8 T cells stimulated with gp33 peptide, (K) splenic CD44^+^ CD8 T cells from Cl13 infected mice treated with ruxolitinib or vehicle at 10 dpi. Statistical comparisons were performed using Fisher’s exact test (B-C), one-way ANOVA with Dunnett’s post-test (D, H), Student’s two-tailed t-test (J, K). Bars represent standard deviation; JAKi, JAK inhibitor; *, *p* ≤ 0.05; **, *p* ≤ 0.01; ***, *p* ≤ 0.001; ****, *p* ≤ 0.0001.

We focused on the JAKi ruxolitinib as it is currently in clinical use for indications including myelofibrosis and graft vs host disease, has a well-established toxicity and pharmacokinetic profile, and was the first JAKi to receive clinical approval (*20*). Ruxolitinib is a dual JAK1 and JAK2 selective inhibitor that exhibits low nanomolar biochemical IC_50_ for both kinases and >100-fold selectivity over JAK3 (*21*). Compound titration revealed a sigmoidal dose response to ruxolitinib in the frequency of YFP-IFN-γ^+^ CD8 T cells (EC50 ∼ 150 nM) (**Figure 1G**). However, the number of YFP-IFN-γ^+^ CD8 T cells showed a bell shape dose response which peaked around 250 nM (**Figures 1G**). Ruxolitinib was superior to aPD-L1 in enhancing the percentage and total number of IFN-γ-producing CD8 T cells as detected by intracellular cytokine staining (**Figure 1D**) as well as increasing IFN-γ protein levels within the supernatants of splenocyte cultures (**Figure 1H**). These results were unexpected given the known immune suppressive effects of JAK inhibition (*22, 23*). Notably, the most dramatically perturbed cytokines in the culture supernatants were myeloid regulators CCL2 and GM-CSF, suggesting there may be a T cell-extrinsic component to ruxolitinib’s effects (**Figure 1I**).

We tested whether ruxolitinib compromised effector functions of CD8 T cells following viral infection *in vivo*. Mice infected with Cl13 and receiving 60 mg/kg daily ruxolitinib exhibited significantly increased total number of GP276-specific splenic CD8 T cells (**Figure S1D**) and of IFN-γ-producing GP276-reactive CD8 T cells than vehicle-treated controls (**Figure 1J**).

Additionally, ruxolitinib reduced the proliferation of naïve T cells *in vitro* stimulated with anti-CD3/CD28, suggesting the enhancement effect is specific to the exhausted T cells (**Figure S1E**). However, viral loads in plasma and organs were unaffected by ruxolitinib treatment at all timepoints examined, suggesting overall CD8 T cell effector function is sufficiently preserved following ruxolitinib treatment early during persistent LCMV infection (**Figure S1F-G**). In contrast, treatment with IFNAR-blocking antibody (aIFNAR) 1 day before Cl13 infection raised viral titres significantly, in line with previous reports, suggesting that ruxolitinib does not fully inhibit IFNAR signaling (**Figure S1F**)(*24, 25*). To substantiate the lack of pro-viral activity of this ruxolitinib dose, we also assessed the frequency of cells positive for viral protein and observed equivalent percentages in ruxolitinib-vs vehicle-treated mice across all immune cell types examined, in contrast to the significantly increased positivity rates in aIFNAR-treated mice (**Figure S1H**). Treatment of ruxolitinib also increased the percentage and total number of proliferative CD8 and CD4 T cells as determined by Ki67 expression (**Figure 1K, S1I**). Ki67^hi^ NK cells were also increased in ruxolitinib treated mice albeit less strikingly than T cells (p ∼ 0.1, **Figure S1I**). Given that ruxolitinib generally exerts antiproliferative effects, we wondered whether the increase in T cells could be partly attributed to reduced cell death. GP33-specific transgenic CD8 T cells (P14) were adoptively transferred to congenic hosts, infected with Cl13 and treated with ruxolitinib or vehicle, then splenic P14 cells analyzed. The percentage of apoptotic P14 cells was reduced more than 3-fold compared to vehicle treated mice (**Figure S1J**). Examining splenocytes of Cl13 infected mice treated with ruxolitinib or vehicle by single-cell RNA-sequencing (scRNAseq) we noticed ruxolitinib increased the percentage of B cells while decreasing the percentage of neutrophils and immature myeloid cells, confirming potential T cell extrinsic effects of ruxolitinib (**Figure S1K**) (*26*). Collectively, these results suggest ruxolitinib treatment can augment cytokine-producing CD8 T cells without compromising viral control, making ruxolitinib an unexpected hit worthy of further investigation.

### Ruxolitinib enhances lymphocyte function and checkpoint blockade efficacy in cancer

We hypothesized that the T cell enhancing effects of ruxolitinib in a persistent viral infection model may be beneficial in enhancing immune checkpoint blockade therapy in cancer. To test this hypothesis, mice received MC38 tumor cells and were treated with two doses of anti-PD1+anti-CTLA4 immune checkpoint inhibitors (ICI) or isotype control 1 week apart as described previously (*27*) and also received ruxolitinib (rux) or vehicle for 7 days between the two immunotherapy doses (**Figure 2A**). In the absence of immunotherapy, ruxolitinib had a marginal effect on tumor growth, however ruxolitinib in combination with ICI (rux+ICI) significantly reduced tumor growth compared to ICI alone (**Figure 2A**). Similar results were observed in the B cell lymphoma model A20 and the lung cancer model LLC1 (**Figure 2B, Figure S2A**). Although it has been shown that ruxolitinib can sensitize anti-PD1-resistant melanoma cells to anti-CTLA4 treatment (*28*), the MC38 and A20 cell lines were not pre-treated with checkpoint inhibitors and do not exhibit prior resistance to anti-PD1.

**Figure 2.**
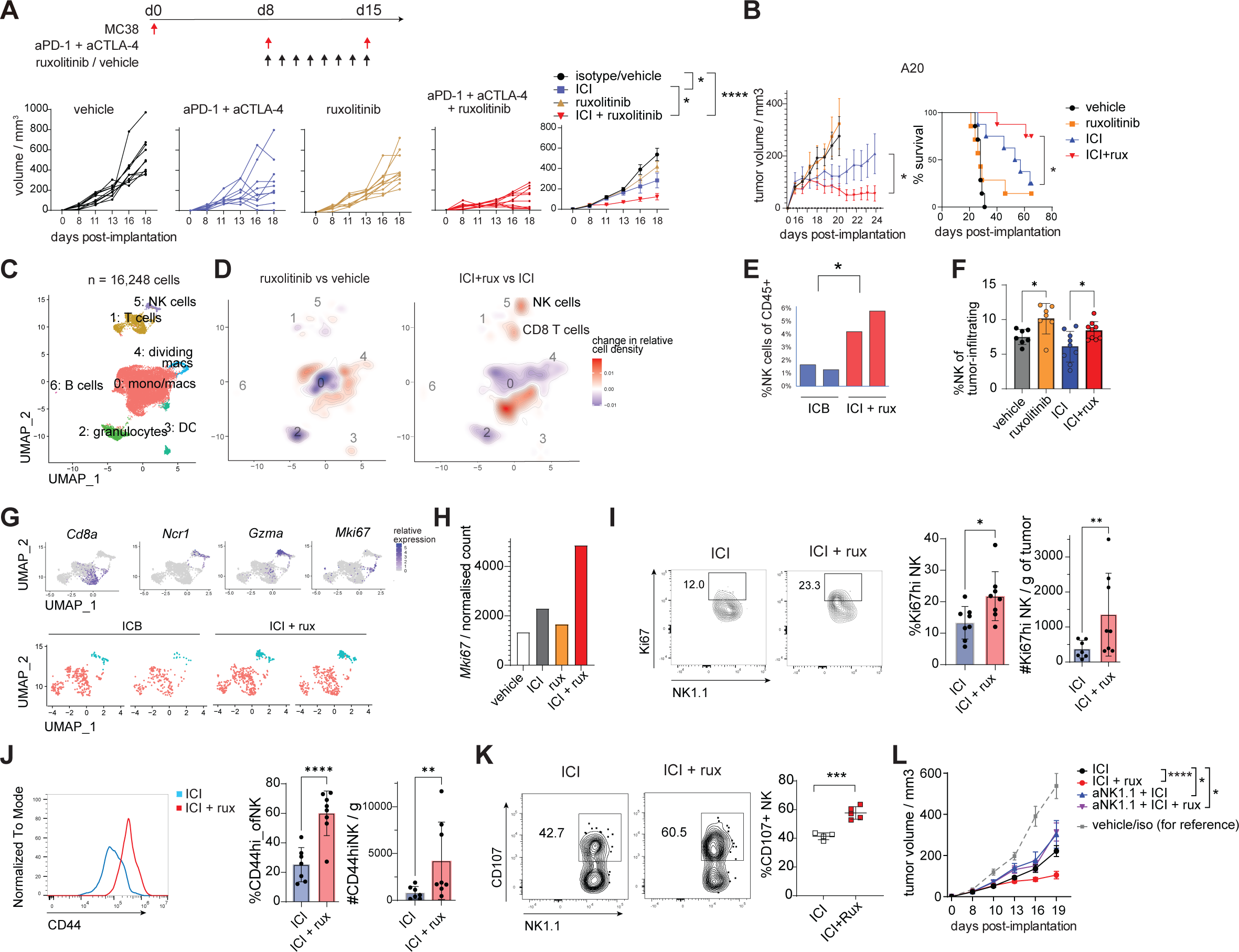
Ruxolitinib enhances checkpoint blockade efficacy and NK cell function. (A, C-K) Mice were implanted with MC38 tumor cells and treated as described in the experimental scheme in 4 experimental groups, tumor measurements were performed every 2-3 days after tumors became palpable; (B) wild type BALB/c mice implanted with A20 tumor cells were treated with ICI or isotype control every 10 days and ruxolitinib or vehicle daily; (C-D) weighted dimensionality reduced map of MC38 tumor-infiltrating cells, treatment groups as described in A; heatmap shows relative protein expression, density plots show relative cell abundance change (normalized to total cell count per sample) in ruxolitinib vs vehicle and ICI+ruxolitinib vs ICI treated mice; (E) percentage of NK cells of MC38 tumor-infiltrating CD45^+^ cells based on scRNAseq clustering; (F) percentage of NK cells of MC38 tumor-infiltrating CD45^+^ cells based on flow cytometry; (G) expression of selected markers in T cell and NK cell scRNAseq clusters; (H) normalized count of *Mki67* in bulk RNA-seq data of sorted tumor-infiltrating NK cells; (I-J) unstimulated tumor-infiltrating NK cells were analyzed by flow cytometry at d18 post implantation; (K) tumor-infiltrating NK cells were treated with PMA/ionomycin and CD107 antibody then analyzed by flow cyometry; (L) mice bearing MC38 tumors were treated with NK cell depleting antibody 2 day before treatment as in panel A, then every 3 days thereafter, tumor sizes measured as described. Statistical comparison of experimental groups was performed by two way-ANOVA (A-B, L), one-way ANOVA and Dunnett’s post-test (F), two-tailed Student’s t-test (E, I-K). Bars represent standard deviation (F, I-J) or s.e.m (A-B, K, L); aPD1, anti-PD1; ICI, immune checkpoint blockade; aNK1.1, anti-NK1.1 depleting antibody; NK, natural killer cell; *, *p* ≤ 0.05; **, *p* ≤ 0.01; ***, *p* ≤ 0.001; ****, *p* ≤ 0.0001.

To understand the basis of this beneficial effect we isolated MC38 tumor-infiltrating cells 1 day following the second dose of anti-PD1+anti-CTLA4 and analyzed single-cell transcriptomes and protein markers using CITE-seq (*29*). Low-resolution clustering identified a prominent cluster of monocytes, macrophages and monocytic dendritic cells, and clusters of T cells, granulocytes, dendritic cells, dividing macrophages, NK cells and B cells (**Figure 2C, S2B-C**). Ruxolitinib treated mice showed major changes in the myeloid compartment and a slight increase in the percentage of NK cells of CD45^+^ cells compared to vehicle treated mice (**Figure 2D**). In contrast, mice receiving ruxolitinib with ICI showed a significantly higher percentage of NK and CD8 T cells of CD45^+^ cells compared to ICI alone (**Figure 2D-E**). This statistically significant increase in NK cells was also observed by flow cytometry and immunohistochemistry (**Figure 2F, S2D**). Tumor-infiltrating NK cells and T cells included a subpopulation expressing *Mki67* and other markers associated with cell division (**Figure 2G**).

Analysis of sorted NK cells by bulk RNA-seq revealed a dramatic upregulation of *Mki67* on tumor-infiltrating NK cells in ICI+rux treated mice compared to ICI alone (**Figure 2H**).

To better define the effect of ruxolitinib on NK cells in ICI-treated tumor-bearing mice, we examined tumor-infiltrating NK cells in the above-described treatment groups by flow cytometry. There was a significant increase in the percentage and total number of Ki67^+^ NK cells in ICI+rux vs ICI treated mice (**Figure 2I**). The expression of CD44 was also increased and the percentage and number of CD44^hi^ NK cells significantly higher in ICI+rux compared to ICI alone (**Figure 2J**). Functionally, we observed significantly increased degranulation of NK cells comparing ICI+rux vs ICI treatment, suggesting ruxolitinib may increase their degranulation capacity (**Figure 2K**).

To determine the causal contribution of CD8 T cells and NK cells to the efficacy of ICI+rux treatment, we depleted CD8 T cells or NK cells immediately prior to treatment with ICI, ruxolitinib or combination therapy and depletion was maintained throughout treatment. As expected, depletion of CD8 T cells in ICI treated mice accelerated tumor growth and the addition of ruxolitinib to ICI did not improve tumor control in CD8 T cell depleted mice (**Figure S2E**). In contrast, NK cell depletion in ICI+rux treated mice mirrored the efficacy observed in mice treated with ICI alone (**Figure 2L, S2F**). This suggests that although CD8 T cells are required for the efficacy of ICI+rux compared to vehicle controls, NK cells are more specifically required for the additional tumor control elicited by ICI+rux compared to ICI alone.

### Ruxolitinib causes reprogramming of granulocytic cells

Our single-cell transcriptomic studies uncovered dramatic changes in tumor-infiltrating myeloid cells induced by ruxolitinib: percentage of granulocytes of tumor-infiltrating CD45^+^ cells was significantly decreased in ruxolitinib-treated and ICI+rux treated mice compared to vehicle and ICI controls, respectively (**Figure 2C-D, 3A**). Flow cytometry confirmed a significant decrease in the percentage and total number of tumor-infiltrating granulocytes in ICI+rux treated mice compared to ICI alone (**Figure 3B**). Granulocytes exhibited significant reductions per gram of tumor both on-treatment and post-treatment in ICI+rux vs ICI mice (**Figure 3B, S3A**). Ruxolitinib also reduced the percentage of splenic neutrophils and the neutrophil-to-lymphocyte ratio in LCMV infected mice, indicating the effect of ruxolitinib on neutrophils is not exclusive to cancer (**Figure S3B-C**).

**Figure 3.**
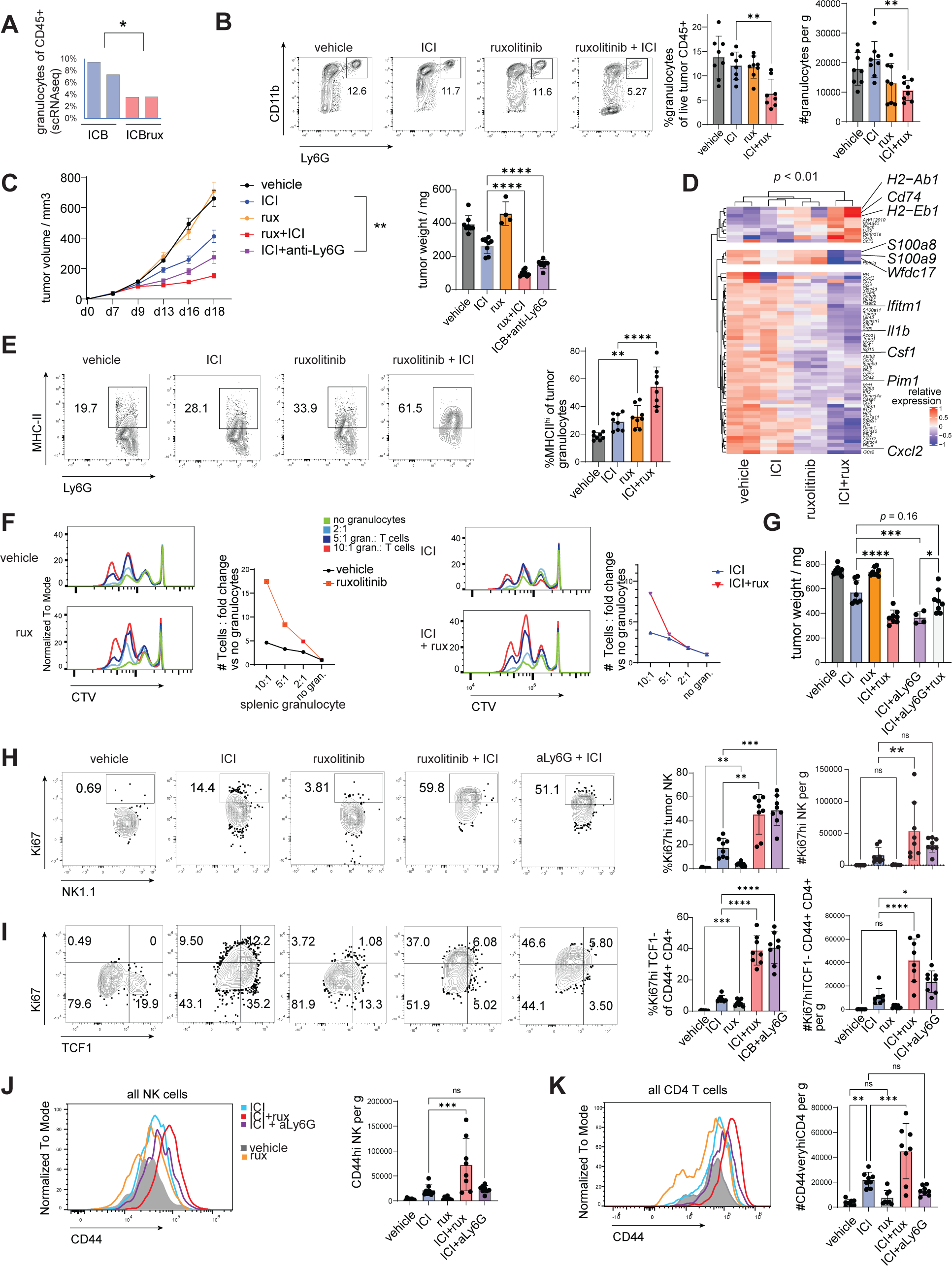
Ruxolitinib reprograms tumor-infiltrating myeloid cells to enhance lymphocyte proliferation. (A) Frequency of granulocytes of CD45^+^ tumor-infiltrating cells based on CITE-seq dataset described in Figure 2C; (B) frequency of granulocytes in MC38 mice treated as described in Figure 2A, flow cytometry performed at d18 post implantation; (C) MC38 tumor bearing mice were treated with Ly6G-depleting antibody or isotype control 2 days prior to treatment with ICI/isotype and ruxolitinib/vehicle, then every 2 days until end of ruxolitinib/vehicle treatment, tumors analyzed at d18 post implantation; (D) genes differentially expressed between ICI+rux treated and ICI treated mice in tumor-infiltrating granulocytes; (E) flow cytometry analysis of tumor-infiltrating granulocytes at d18 post implantation, treatment groups as described in Figure 2A; (F) Splenic granulocytes (CD11b^+^ Ly6G^hi^ Ly6C^int^) from mice treated as described in Fig. 2A were sorted and mixed with purified, CTV-labeled T cells at the ratios indicated and cell mixtures subjected to anti-CD3/CD28 stimulation, flow cytometry analysis at d3 of assay; (G) MC38 tumor bearing mice were treated as described in Figure 3C with the additional experimental group treated with anti-Ly6G, ICI and ruxolitinib, tumors analyzed 2d after treatment cessation; (H-K) MC38 tumor bearing mice were treated as in Figure 3C and tumor-infiltrating cells analyzed 2d after treatment cessation. Statistical comparison of treatment groups was performed using two-tailed Student’s t-test (A), one-way ANOVA with Dunnett’s post-test (B, C right panel, E, G-K), two-way ANOVA (C left panel) or DESeq2 Wald test (D); ICI, immune checkpoint blockade; CTV, cell trace violet; *, *p* ≤ 0.05; **, *p* ≤ 0.01; ***, *p* ≤ 0.001; ****, *p* ≤ 0.0001.

Granulocytic cells include a population of polymorphonuclear myeloid derived suppressor cells (PMN-MDSCs) which share surface markers with neutrophils (*30*). To assess the contribution of granulocytic myeloid cells to ruxolitinib’s enhancement of ICI efficacy, Ly6G^+^ cells were depleted using an anti-Ly6G depleting antibody (aLy6G) prior to ICI, rux or ICI+rux treatment and depletion was maintained throughout the treatment period. Interestingly, depletion of Ly6G^+^ cells in ICI-treated mice significantly reduced tumor growth and tumor weight, confirming the negative contribution of these cells to antitumor immune response in this model (**Figure 3C**). To directly demonstrate that MC38 tumor-infiltrating myeloid cells are suppressive, the cells were subjected to a suppression assay: Gr1^+^ cells were isolated from MC38 tumor bearing mice and tested for their ability to suppress IL-2-driven proliferation and IFN-g production of NK cells. As expected, increasing doses of Gr1^+^ cells reduced the percentage and total number of IFNg^+^ NK cells, confirming their suppressive abilities (**Figure S3D**). Consistently, splenic granulocytes expressed arginase in MC38 tumor-bearing mice and ruxolitinib nearly ablated arginase 1 (ARG1) protein levels in granulocytes both when administered alone or in conjunction with ICI, suggesting ruxolitinib not only modulates granulocyte numbers but also the expression of suppressive markers (**Figure S3E**). Moreover, tumor-infiltrating granulocytes in ICI+rux vs ICI treated mice showed significantly reduced expression of MDSC-associated markers *Csf1*, *Cxcl2*, *Ifitm1*, *Il1b*, *S100a8*, *S100a9* and *Wfdc17* (**Figure 3D, S3F**). The tumor infiltrating neutrophils exhibited higher expression of MHC class II molecules as well as *Cd74* in ICI+rux treated mice, suggesting they acquired transcriptomic features of immune stimulatory antigen presenting cells (**Figure 3D-E, S3F**). Moreover, the induction of MHC-II expression in granulocytes was also observed in ruxolitinib-treated mice infected with Cl13, indicating the effect is not limited to cancer (**Figure S3G**). To explore the functional relevance of these changes, we performed an *in vitro* T cell proliferation assay using splenic granulocytes from MC38 tumor bearing mice treated with vehicle, ruxolitinib, ICI or ICI+rux. Remarkably, granulocytes from ruxolitinib treated mice enhanced T cell division and the total number of T cells over 3 days of stimulation with anti-CD3/CD28 to a significantly greater extent than ICI or vehicle derived granulocytes (**Figure 3F**).

To determine the contribution of these granulocytes to the synergy between ICI and ruxolitinib, we depleted Ly6G^+^ cells during ICI/ICI+rux treatment and assessed tumor growth. Whereas rux+ICI and anti-Ly6G+ICI (aLy6G+ICI) groups showed significantly reduced tumor weight compared to ICI alone, ruxolitinib provided no additional benefit to aLy6G+ICI treated mice compared to rux+ICI or aLy6G+ICI groups (**Figure 3G**).

To identify which antitumor immune responses were invigorated by granulocyte depletion or by ruxolitinib treatment, tumor-infiltrating cells were investigated by flow cytometry after treatment cessation. The effect of aLy6G+ICI on Ki67 NK cells phenocopied the effect of rux+ICI with a >2 fold increase in the percentage of Ki67^+^ NK cells in these groups compared to ICI alone (**Figure 3H**). A similar effect was observed on CD4 T cells, with a particularly striking increase in TCF1^-^ CD44^hi^ CD4 T cells in both ICI+rux and ICI+aLy6G groups (**Figure 3I**). In contrast, the increased expression of CD44 on NK cells was specific to the ICI+rux group, suggesting ruxolitinib also has granulocyte-independent effects (**Figure 3J**). The expression levels of CD44 on CD8 T cells was also enhanced in ICI+rux compared to other groups but the total number of CD44^+^ CD8 T cells was not significantly increased by ICI+rux or ICI+aLy6G compared to ICI (**Figure S3H**). In summary, ruxolitinib treatment in combination with ICI results in altered transcriptional profile and reduced numbers of Ly6G^+^ cells in the tumor and spleens of tumor bearing mice, and Ly6G^+^ cells play a causal role in the ruxolitinib-mediated increase in Ki67^+^ NK cells and enhancement of ICI efficacy.

### Ruxolitinib reduces suppressive gene expression in myeloid cells of the bone marrow, blood and tumor

We asked whether the effect of ruxolitinib on tumor-infiltrating myeloid cells had conserved patterns across different tumor types. To address this, CITE-seq data of tumor-infiltrating myeloid cells from 3 different mouse tumor types treated with ICI+rux or ICI were integrated and subjected to clustering. High diversity was found in the monocytic/macrophage cluster whereas only one granulocyte cluster emerged at that resolution (**Figure 4A-B, S4A**). In a striking concordance between tumor types, the percentage of myeloid cells in Clusters 0 (ARG1^+^ mac/mono cells) and 3 (granulocytic cluster) was reduced in ICI+rux compared to ICI treated mice (**Figure 4C-D**). The loss of MDSC markers and gain of MHC gene expression by granulocytes was observed in all 3 tumor types with ICI+rux treatment compared to ICI treated controls (**Figure S4B**). In contrast, cells in Cluster 2 expressing markers of monocyte derived dendritic cells (moDC) increased in frequency in all 3 tumor types and the ISG-expressing moDC Cluster 8 increased in the LLC1 and MC38 models (**Figure 4D**). To further understand the phenotypic characteristics of these clusters, we compared markers of human monocytes and macrophages in the MoMac-VERSE (*31*). Cluster 0 expressed multiple markers such as *Spp1*, *Fabp5* and *Apoe* which corresponded to the TREM2^+^ macrophage #3 cluster of the MoMac-VERSE, the cluster most significantly and consistently increased in human cancers compared to non-diseased tissue of all MoMac-VERSE clusters (**Figure 4D, S4C**) (*31*).

**Figure 4.**
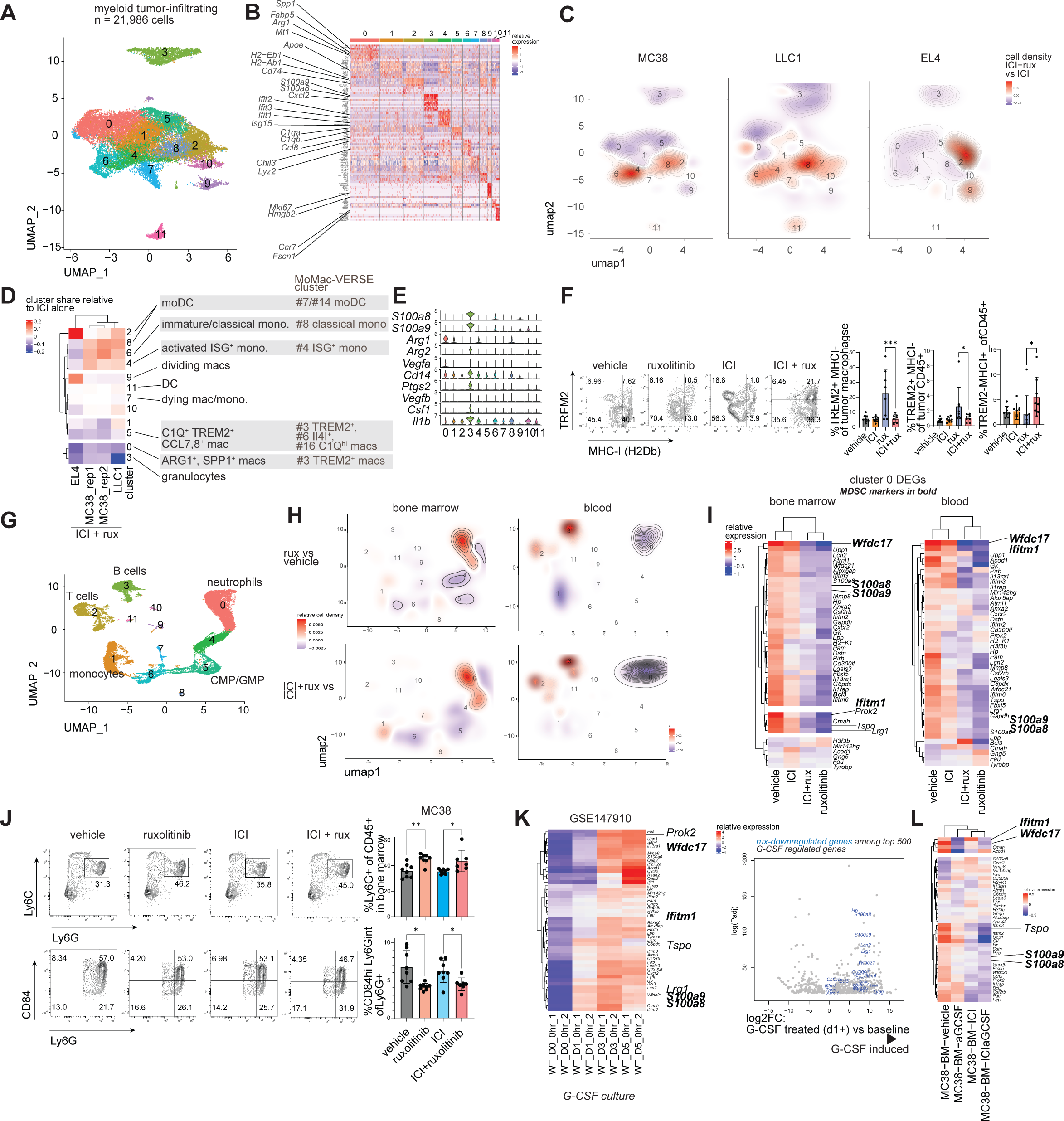
Myeloid reprogramming by ruxolitinib across tumor types and organ sites. (A-E) Integrated CITE-seq analysis of tumor-infiltrating myeloid cells from EL4, LLC1 and MC38 tumors: (A) dimensionality-reduced plot and clustering of integrated dataset; (B) mRNA markers of 11 clusters; (C) difference in relative cell density in ICI+rux treated mice compared to respective ICI treated controls; (D) difference in relative cluster share between ICI+rux treated mice and respective ICI treated controls, right: MoMac-VERSE (*31*) clusters most closely aligned with the observed clusters based on marker genes; (F) tumor-infiltrating macrophages in mice treated as in Figure 2A were analyzed by flow cytometry 2 days after treatment; (G-I) bone marrow and blood from mice treated as in Figure 2A were analyzed by CITE-seq: (G) dimensionality reduced map and clustering of combined dataset; (H) difference in relative cell density in ruxolitinib vs vehicle treated mice and ICI+rux vs ICI treated mice; (I) relative expression of differentially expressed genes between ruxolitinib treated and vehicle treated mice in cluster 0 (differentiated neutrophils), MDSC markers shown in bold italic; (J) flow cytometric analysis of bone marrow granulocytes in mice treated as in Figure 2A analyzed 2 days after treatment; (K) expression of ruxolitinib-regulated genes in myeloid cells cultured in the presence of G-CSF for 0-5 days (GSE147910), MDSC markers in bold italic, right: volcano plot of G-CSF regulated genes at d1-5 vs d0, ruxolitinib-inhibited genes highlighted in blue; (L) MC38 tumor bearing mice were treated with low-dose G-CSF neutralizing antibody or isotype control every 3 days once palpable and ICI or isotype control once palpable and 7 days later, bone marrow granulocytes analyzed by CITE-seq 2 days after end of treatment, heatmap shows relative expression of ruxolitinib-downregulated genes; DEG, differentially expressed gene; moDC, monocyte-derived dendritic cell, mac, macrophage; ICI, immune checkpoint blockade. Statistical comparison of experimental groups was performed using one-way ANOVA with Dunnett’s post-test (F, J) or DESeq2 Wald test (I, K); *, *p* ≤ 0.05; **, *p* ≤ 0.01; ***, *p* ≤ 0.001.

Furthermore, Cluster 0 and 3 expressed the highest levels of MDSC markers of all myeloid clusters (**Figure 4E**). These observations suggested ruxolitinib causes a relative reduction in myeloid cells expressing suppressive markers while increasing the relative abundance of MHC molecule-expressing moDCs and granulocytes. The increase of MHC-II-expressing Ly6C^hi^ monocytic cells was also observed by flow cytometry (**Figure S4D**). Flow cytometry also confirmed a switch from TREM2^+^ MHC^-^ macrophages to TREM2^-^ MHC^+^ in ICI+rux compared to ICI (**Figure 4F**). Furthermore, ruxolitinib treatment also increased MHC-II expression and reduced *S100a8* expression in splenic monocytes, and switch from TREM2^+^ MHC^-^ to TREM2^-^ MHC^+^ macrophages in the persistent viral infection model (**Figure S4E-F**). These data reveal a broad ruxolitinib-induced shift from suppressive gene expression to MHC expression in granulocytic, monocytic and macrophage populations.

Suppressive activity in myeloid cells can be acquired due to excessive stimulation by soluble factors including the JAK-signaling cytokines G-CSF and GM-CSF (*9, 32, 33*). We asked whether ruxolitinib interfered with the suppressive programming of myeloid cells in the tumor and/or during their development in the bone marrow. To address this question, tumor bearing mice were treated as described in Figure 2A and their peripheral blood and bone marrow CD45^+^ cells were analyzed by single-cell transcriptome sequencing. As expected, ruxolitinib and ICI+rux treated mice showed a reduction in blood neutrophils and increase in lymphocytes compared to vehicle and ICI treated mice, respectively (**Figure 4G-H**). Blood monocytes from ICI+rux treated mice expressed lower levels of suppressive markers *Arg2*, *S100a8* and *S100a9* (**Figure S4G**). There was a significant increase in the percentage of differentiated neutrophils in the bone marrow of ruxolitinib and ICI+rux treated mice compared to respective controls, whereas immature neutrophils were underrepresented in these mice based on Neutrotime gene classification (**Figure 4G-H, S4H**) (*34*). Analysis of differentially expressed genes revealed the downregulation of multiple previously identified MDSC markers in the neutrophils of ruxolitinib and ICI+rux treated mice including *Wfdc17*, *S100a8*, *S100a9* and *Ifitm1* (**Figure 4I**) (*9, 35*). These MDSC markers were reduced in both bone marrow and blood neutrophils, suggesting that ruxolitinib may influence the gene expression of neutrophils before they leave the bone marrow. There was a dramatic reduction in the expression of myeloid migration genes *Lrg1*, *Prok2* and *Tspo* in neutrophils of ruxolitinib treated mice (**Figure 4I**) (*36-39*). t However, ruxolitinib did not significantly reduce the transcriptomic program associated with normal neutrophil differentiation as defined by high Neutrotime score, suggesting ruxolitinib does not prevent neutrophil differentiation (**Figure S4I**) (*34*).

To validate the scRNAseq-identified changes, we examined bone marrow neutrophils by flow cytometry and observed significant increase in the percentage of neutrophils in ruxolitinib treated mice compared to controls (**Figure 4J**) The PMN-MDSC protein marker CD84 was significantly downregulated while the total number of CD84^-^ neutrophils did not decrease (**Figure 4J, S4J**). Importantly, the percentage of CD84^+^ BM neutrophils was also reduced in other tumor models including the T cell lymphoma EL4, indicating these effects are not unique to MC38 or colorectal cancer (**Figure 4J, S4K**).

To identify upstream signals targeted by ruxolitinib which lead to the observed gene expression changes, we analyzed the predicted cytokine activity in BM neutrophils using Cytosig (**Figure S4L**) (*40*). Several cytokines showed dominant activity: EGF, IL-17A, IL-36, NO, G-CSF and TRAIL. Both IL-17 and G-CSF regulate emergency granulopoiesis and suppressive myeloid development, IL-36 regulates neutrophilic inflammation and NO mediated the suppressive activity of MDSCs. Of these, only G-CSF uses canonical JAK1 or JAK2 signaling. To determine whether G-CSF activity played a role in inducing ruxolitinib-downregulated genes, the temporal effect of G-CSF on myeloid cell transcriptomes was analyzed. Strikingly, virtually all ruxolitinib-downregulated genes were induced by G-CSF in the publicly available datasets examined, including genes such as *Wfdc21*, *Wfdc17* and *Ifitm6* which were among the most highly induced genes by G-CSF based on fold change and p value (**Figure 4K**). We treated MC38 tumor-bearing mice with the G-CSF neutralizing antibody (aG-CSF) with or without ICI. aGCSF treatment caused reduced expression of multiple G-CSF targets including *Wfdc17* in bone marrow neutrophils, indicating that G-CSF is required for the observed expression levels of these genes in vehicle- and ICI-treated MC38 tumor bearing mice (**Figure 4L**).

Given that ruxolitinib has been reported to modulate cancer cell-intrinsic resistance to anti-PD1, we wondered whether cancer cell-intrinsic JAK1 or JAK2 were required for the effects observed above. To address this, JAK1 and JAK2 deficient MC38 cells generated previously (*41*) were injected in B6 hosts and treated as described in Fig. 2A, then tumor infiltrating cells analyzed by flow cytometry. Although growth of *Jak1*^-/-^ MC38 was impaired *in vivo* as previously reported (*41*), we observed an increase in the percentage of monocyte-derived dendritic cells in tumors of ICI+rux vs ICI treated mice by single-cell transcriptomics (**Figure S4M**). Monocytes also expressed significantly higher transcript and protein levels of MHC-II in ICI+rux vs ICI treated tumors (**Figure S4N-O**). Similar to wt MC38, tumor-infiltrating macrophages in *Jak1*^-/-^ MC38 tumors exhibited lower expression of TREM2 and higher expression of MHC-II (**Figure S4M, P).** We also observed significantly higher expression of MHC-II in both granulocytic and monocytic tumor-infiltrating populations in *Jak2*^-/-^ MC38 tumors (**Figure S4Q**). In agreement with the effects of ruxolitinib in the LCMV Cl13 model, these data demonstrate that although JAK1/2 deficiency alters the tumor-immune properties of syngeneic tumors, ruxolitinib’s myeloid reprogramming effect is at least partially independent of tumor cell-intrinsic JAK1 and JAK2.

### Blood changes associated with nivolumab response in ruxolitinib-treated patients with resistant or refractory Hodgkin lymphoma

To investigate whether ruxolitinib can enhance antitumor immune responses in humans, we studied a clinical trial of ruxolitinib with nivolumab in Hodgkin lymphoma patients who previously failed checkpoint inhibitor therapy. In a Phase 1/2 open-label trial, eligible patients received ruxolitinib orally (dose ranged from 5-20 mg twice daily) followed by nivolumab (480 mg intravenously every 4 weeks) (NCT03681561). We examined peripheral blood samples collected pre-treatment and 1 week after the start of ruxolitinib treatment, allowing for the effect of ruxolitinib to be evaluated before nivolumab was given. Of the 19 patients treated to date, there were 14 complete paired blood samples pre and post ruxolitinib available for analysis.

Bulk RNA-sequencing of peripheral mononuclear cells identified 155 genes differentially expressed post-ruxolitinib vs baseline, including the most prominently reduced genes *CCL2* and *CCL7* (**Figure 5A**). A computational deconvolution of cell types using a large reference single-cell CITE-seq PBMC dataset revealed that lymphocytes increased in percentage accompanied by a sharp decrease in myeloid progenitors and monocytes (**Figure 5B, S5B-C**). Concordantly, absolute lymphocyte counts determined by hematology analyzer significantly increased following ruxolitinib treatment (**Figure 5C**). Mapping differentially expressed genes onto the PBMC reference dataset showed that whereas ruxolitinib-upregulated genes were distributed relatively evenly across B cells, NK cells and CD8 T cells, ruxolitinib-downregulated genes were particularly enriched in CD14^+^ monocytes (**Figure S5D**). The recently published cohort of nivolumab-treated Hodgkin lymphoma patients indicated that B cells and NK were higher at baseline in nivolumab responders than in non-responders, suggesting that ruxolitinib preferentially enhances cells types associated with response to nivolumab (*42*). Interestingly, the percentage of cytokine producing CD8 T cells was unimpaired and the absolute count of cytokine-producing CD8 T cells significantly increased post-treatment, suggesting ruxolitinib does not adversely affect CD8 T cells in Hodgkin lymphoma patients treated in this trial (**Figure 5D**).

**Figure 5.**
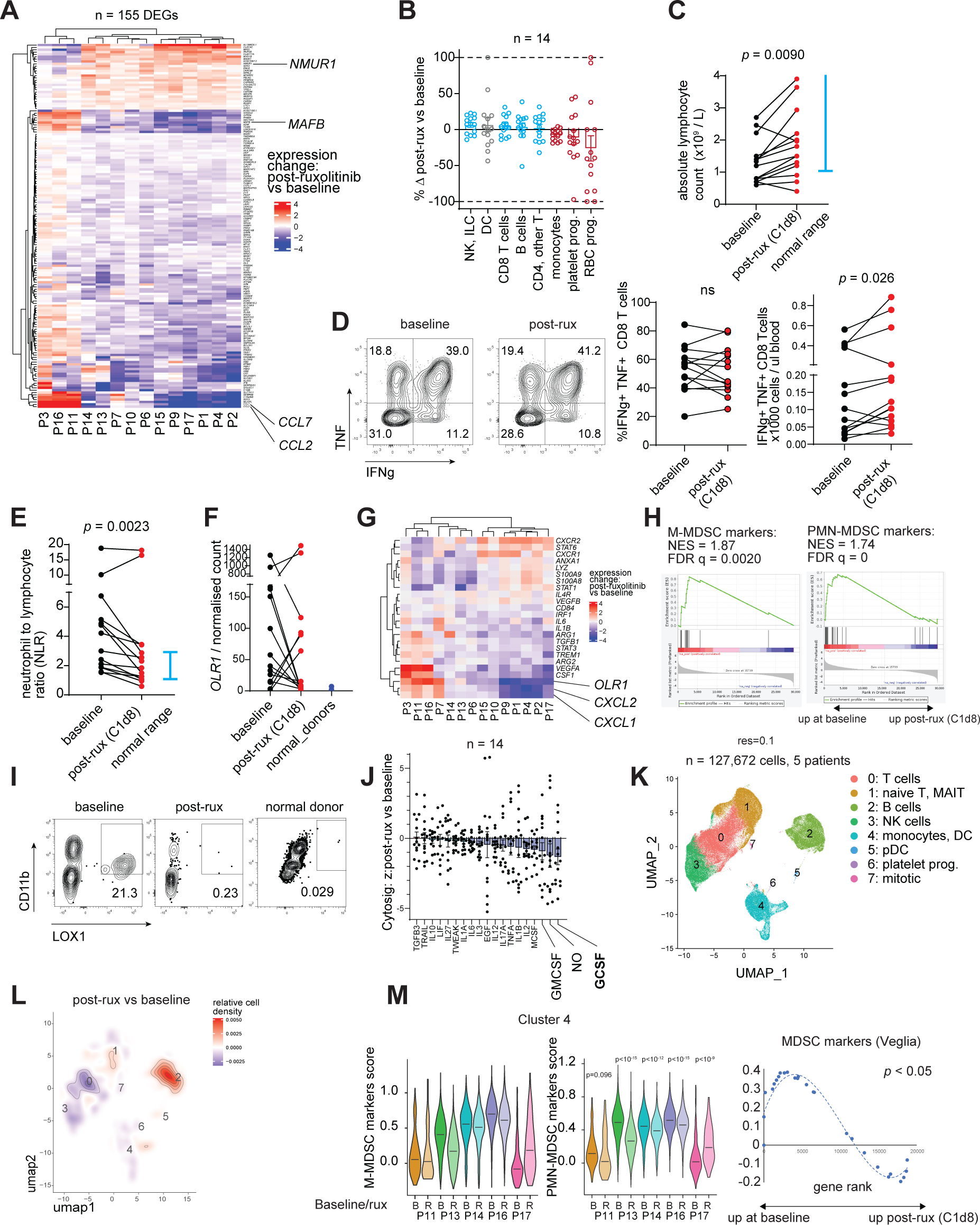
Ruxolitinib-induced peripheral blood changes in patients with anti-PD1-resistant or refractory classical Hodgkin lymphoma. Peripheral blood samples were collected at baseline and 8 days post-ruxolitinib treatment (C1d8), subjected to hematologic analysis and mononuclear cells isolated for further analysis: (A) PBMC bulk transcriptome changes plotted as post-ruxolitinib vs baseline, only significantly differentially expressed genes shown; (B) changes in cell type percentages post-ruxolitinib vs baseline as predicted from bulk transcriptomes; (C) absolute lymphocyte counts established by hematologic analyzer; (D) PBMCs were stimulated with PMA/ionomycin for 3h and CD8 T cell cytokines examined by flow cytometry; (E) neutrophil-to-lymphocyte ratio determined by hematologic analyzer; (F) expression of *OLR1* in bulk RNA-seq from patient and normal donor PBMCs; (G-H) relative expression of MDSC marker genes in PBMCs after ruxolitinib treatment compared to baseline; (H) GSEA of M-MDSC and PMN-MDSC genesets from Veglia et al.; (I) flow cytometric detection of LOX1^+^ PMN-MDSCs in patients but not normal donors; (J) relative change in cytokine transcriptomic score after ruxolitinib compared to baseline, MDSC associated cytokines highlighted; (K-M) single-cell RNA-seq analysis of live PBMCs from 5 patients pre- and post-ruxolitinib: (K) integrated dimensionality reduction and clustering and relative change in cell frequency post-ruxolitinib; (L) relative cell density in post-ruxolitinib samples compared to baseline, aggregated across patients; (M) MDSC signature score expression before and after ruxolitinib, GSEA plot shows change in combined MDSC marker geneset; DEG, differentially expressed gene; GSEA, gene set enrichment analysis; NES, normalized enrichment score; DC, dendritic cell; NK, natural killer cell; RBC, red blood cell; prog., progenitor cell; C1d8, Cycle 1 day 8; FDR, false discovery rate; MAIT, mucosal associated invariant T cell. Statistical comparison of experimental groups was performed using DESeq2 Wald test (A), paired Student’s t-test (C, D, E) or GSEA test (H, M) (*84*).

The neutrophil-to-lymphocyte ratio, clinically known to correlate with MDSC frequency and poor prognosis across cancer types including Hodgkin lymphoma (*43-45*), showed a significant reduction post-ruxolitinib (**Figure 5E**). Assessing the expression of published MDSC markers (*9*) revealed a statistically significant reduction of their expression after ruxolitinib, with particularly striking decreases for PMN-MDSC markers *OLR1*, *CXCL1* and *CXCL2* (**Figure 5F-H**). Gene expression of *OLR1* correlated with elevated protein expression analyzed by flow cytometry and no expression of *OLR1* was detected in normal donor PBMCs (**Figure 5I**). The expression of G-CSF target genes was significantly downregulated after ruxolitinib treatment (**Figure 5J**) including certain markers upregulated in G-CSF mobilized patient monocytes compared to normal CD14^+^ monocytes such as *ADM* (**Figure S5E**). Transcription factors STAT1, STAT3 and STAT6 play a role in MDSC programming and signal downstream of JAK1 and JAK2 (*9*). Of these, STAT3 signals downstream of G-CSF. Gene set enrichment analysis of annotated STAT targets showed a significant reduction in STAT3 target genes post ruxolitinib (**Figure S5F**).

To obtain independent verification of the effects of ruxolitinib on PBMCs, a representative subset of patients was subjected to single-cell RNA-seq analysis of PBMCs at baseline and post-ruxolitinib. Following quality control and inter-subject integration, 127,672 cells separated into clusters associated with distinct cell types (**Figure 5K**). The relative proportion of pDCs, B cells and naïve T cells increased in most patients after ruxolitinib in agreement with bulk RNA-seq and complete cell count data (**Figure 5L, S5G**). In contrast, the percentage of classical monocytes of total PBMCs decreased significantly in most patients as suggested by bulk RNA-seq (**Figure 5L, S5G**). At the transcriptome level, the monocytic cluster showed a statistically significant reduction in MDSC marker genes (**Figure 5M**). The protein marker CD84 showed a modest but consistent reduction in the monocytes of all example patients examined by flow cytometry (**Figure S5H**). These data demonstrate that MDSC marker reductions in the blood of Hodgkin lymphoma patients treated with ruxolitinib are not only caused by relative reductions in myeloid cells but also by reduced expression of MDSC markers in monocytic cells.

Although this clinical trial is still in progress, we observed preliminary evidence of clinical efficacy: interim trial data reported a best overall response rate of 75% among 16 evaluable patients including 3 complete and 2 partial responders (*46*). This is higher than published response rates for ruxolitinib monotherapy (9 and 54%) and nivolumab monotherapy (69%) in patients naïve to respective agents (*47-49*). Taken together, these data suggest ruxolitinib treatment in checkpoint inhibitor non-responsive patients causes a partial normalization in blood cell composition consistent with depletion of MDSCs and could be useful in enhancing response to anti-PD1 therapy.

## Discussion

Myeloid suppressor cells are an increasingly appreciated factor influencing the response to cancer immunotherapy, yet their clinical targeting remains to be realized. JAK inhibitors target pathways directly involved in the suppressive programming of MDSCs, however the JAK inhibitors AZD1480 and tofacitinib were reported to expand MDSCs or increase their suppressive capacity in some disease models (*12, 13*). Moreover, JAK inhibitors exert direct effects on cancer cells (*28*), dendritic cells (*50*), NK cells (*51*), T cells (*52*) and other cell types (*26*), suggesting myeloid cells are influenced by both direct and indirect effects of JAK inhibition. Our results suggest that in the presence of checkpoint inhibitors, ruxolitinib inhibits suppressive gene expression in granulocytic, monocytic and macrophage cells while increasing their MHC expression. It is possible that these effects are dependent on the inhibitor dose as well as the inhibitor’s JAK selectivity and pharmacological profile. Given the important role of JAK-STAT signaling in antigen presentation and T cell recruitment, complete ablation of JAK1/2 activity is unlikely to improve antitumor immunity or ICI response (*41, 53*). Indeed, ruxolitinib failed to increase the total number of cytokine-producing CD8 T cells *in vitro* above 250 nM. However, at the doses tested, T cell function and total numbers were preserved in mice *in vivo* as well as Hodgkin lymphoma patients. This is echoed by observations in other studies in which T cell function was preserved at therapeutic doses of JAK inhibitors (*54, 55*). Therefore, drug dosing is likely of key importance to the success of JAK inhibitors in the context of cancer immunotherapy.

Classic Hodgkin lymphomas exhibit frequent *JAK2* copy number changes (*56-58*). Ruxolitinib previously showed some efficacy as a monotherapy in Hodgkin lymphoma: in a phase II trial, ruxolitinib monotherapy showed 9.4% (3/32) overall response rate with a median response of 7.7 months (*47*); Kim et al. observed a best response rate of 54% (7 of 13 patients had complete, partial response or stable disease) with a median response time of 5.6 months (*48*). Patients in these trials were not refractory or resistant to nivolumab. In our trial, interim analysis shows a best overall response rate of 75% with a median of 12.5 months in a checkpoint inhibitor relapsed or refractory cohort. This result echoes anecdotal observations of responses in Merkel cell carcinoma and non-Hodgkin B cell lymphoma in which patients on JAK inhibitors responded to anti-PD1 treatment and suggests the combination of ruxolitinib with nivolumab may be more efficacious than either therapy alone (*59, 60*). The preliminary clinical experience offers the possibility that the antitumor immune response may be enhanced by ruxolitinib in this patient population, as supported by the peripheral blood decrease in MDSC markers after ruxolitinib treatment. As ruxolitinib caused comparable patterns of myeloid reprogramming in the LCMV C13 model, multiple syngeneic cancer models and in mice bearing *Jak1*^-/-^ and *Jak2*^-/-^ MC38 tumors, it is likely that at least some of its immune-mediating effects are independent of cancer cell-intrinsic JAK signaling.

Previously, ruxolitinib was shown to reduce recruitment of inflammatory neutrophils to inflamed tissues in a cytokine-dependent manner in the bHLH model (*61*). Given that MDSCs are thought to originate through inappropriate myeloid activation including high levels of inflammatory cytokines such as IL-1β and IL-6, it is possible that some molecular mechanisms are shared between the recruitment of inflammatory neutrophils in this disease model and the recruitment of suppressive granulocytic MDSCs in cancer (*8, 9*). LRG1 regulates neutrophil chemotaxis in a STAT3-dependent manner (*36*). TSPO modulates neutrophil migration and adhesion (*37*). PROK2/Bv8 enhances mobilization of Gr1^hi^ myeloid cells from the bone marrow and is an investigational target for cancer therapy (*38, 39*). The robust effect of ruxolitinib on the expression of chemokines and chemotactic genes in granulocytes in the bone marrow, blood and tumor suggest that ruxolitinib modulates general neutrophil trafficking in addition to their acquisition of a suppressive transcriptomic profile.

Given the occurrence of suppressive myeloid cells in various tumor types (*62*), it is possible that JAK inhibition could improve checkpoint responses in other malignancies. Hodgkin lymphoma may be a particularly good target for ruxolitinib treatment because of the known myeloid expansion confirmed by negative prognostic relevance of NLR for PFS (*45, 63*) and negative prognostic relevance of monocytes for nivolumab response. However, NLR is robustly associated with anti-PD1 response in other tumor types including melanoma and lung cancer (*64, 65*). Therefore, our data imply that ruxolitinib could be synergistic with checkpoint blockade in cancer therapy and this therapeutic combination should be explored in future well-designed clinical trials.

## Materials and methods

### Experimental design

Analysis of blood samples collected in an ongoing clinical trial was performed using the samples available without a prespecified target sample size. Sample size for preclinical tumor studies was selected to detect an effect size of at least 1.5 as measured by tumor weight.

### Human samples

We analyzed blood samples from patients with Hodgkin lymphoma treated in an ongoing clinical trial of ruxolitinib with nivolumab (NCT03681561). Pre-planned blood collection for correlative endpoints occurred at baseline, 1 week of ruxolitinib therapy and 2 weeks of combined therapy of ruxolitinib with nivolumab. All patients signed IRB-approved informed consent for treatment and sample collection and were treated in accordance with the Declaration of Helsinki.

Peripheral blood mononuclear cells were isolated and frozen according to standard protocols. Normal peripheral blood mononuclear cells were obtained through the TSRI Normal Blood Donor Services program (TSRI IRB #15-6710).

### Animal studies

Mouse studies were approved by the Institutional Animal Care and Use Committee of The Scripps Research Institute (TSRI IACUC #15-0017) and performed in accordance with its guidelines. P14 mice were generated by Pircher et al. and are available as Jackson Laboratory stock 37394 (*66*).

### Virus

LCMV Clone 13 was passaged on BHK cells as previously reported (*67*). Serum was isolated by low-speed centrifugation to remove red and white blood cells. 10 µl of serum was used to perform 10-fold serial dilutions and quantified by focus forming assay on VeroE6 cells as previously described (*68*).

### Kinase inhibitor library screen

A library of kinase inhibitors was obtained from a commercial source (cat. HY-L008, MedChemExpress, Monmouth Junction, NJ). Spleens from LCMV-CL13 infected IFN-γ-YFP mice were isolated at day 15 p.i. and single cell suspensions prepared. Following depletion of B cells, 2x10^5^ cells were seeded in duplicate onto 96-well plate wells pre-spotted with DMSO or library compounds at 1000 nM, 500 nM, 250 nM and 100 nM final concentration. Following a 5-day incubation period, the viability dye 7-AAD was added to each well (1:50 dilution) and plates were rested for 15 minutes. Cells were then analyzed on a ZE5 flow cytometer (Bio-Rad). Z scores were calculated for the frequency of YFP^+^ 7AAD^-^ cells in compound wells compared to DMSO control wells.

### Compound target gene enrichment analysis

Plate list of compounds in the ReFrame library (*69*) was filtered for duplicates such that only a single plate well per unique molecule was included. Compounds with autofluorescence in the YFP channel were removed by applying the threshold raw %YFP^-^ CD8 T cells > 25. To obtain drug gene interactions, ChEMBL IDs were first retrieved using the ChEMBL API (Plate list of compounds in the ReFrame library (*69*) was filtered for duplicates such that only a single plate well per unique molecule was included. Compounds with autofluorescence in the YFP channel were removed by applying the threshold raw %YFP^-^ CD8 T cells > 25. To obtain drug gene interactions, ChEMBL IDs were first retrieved using the ChEMBL API (https://chembl.gitbook.io/chembl-interface-documentation/web-services/chembl-data-web-services), then the Drug Gene Interaction Database queried using ChEMBL IDs (*18*). JAK inhibitors were defined as compounds with a curated interaction with JAK1, JAK2 or JAK3. Fisher’s exact test was used to assess enrichment of JAK inhibitors among hit compounds.

### T cell proliferation and intracellular cytokine detection assays

Splenocytes from LCMV-Cl13 infected IFN-γ-YFP mice were labeled with CellTrace Violet (CTV) and seeded onto round-bottom 96-well plates at 2x10^5^ cells/well in complete T cell media (10% FBS, L-glutamine, Pen/Strep, NEAA, Sodium Pyruvate, HEPES, β-mercaptoethanol) supplemented with a cocktail of all known immunodominant LCMV CD8-specific (LCMV-GP33-41, LCMV-GP276-286 and LCMV-NP396-404) and CD4-specific (LCMV-GP61-80) epitope peptides (*70, 71*). Cells were treated with either DMSO, 250 nM ruxolitinib phosphate (LC Laboratories, Woburn, MA; Cat R-6688), 50 µg/ml anti-IFN-γ (BioXCell, Cat # BE0055) or 25 µg/ml anti-PD-L1 (BioXCell, Cat # BE0101). Brefeldin A (Thermo Fisher, Cat#: B7450) was added at a 1:500 dilution to each well 6 h prior to cell isolation to block T cell-specific cytokine secretion. Surface antigens were stained for 30 minutes on ice, cell fixed and permeabilized using the Transcription Factor Staining Buffer Set (Cat #: 00-5523-00). Permeabilized cells were then stained for intracellular antigens. To track GP-specific CD8 T cells in vitro, 1,000 congenic Thy1.1^+^ CD8^+^ T cells (P14) from TCR tg mice that recognize the LCMV-GP33-41 epitope (Pircher et al. 1989) were engrafted into IFN-γ-YFP mice 1 day prior to LCMV infection.

### Extracellular cytokine detection assay

Supernatants from in vitro IFN-γ-YFP cultures were isolated at day 3 and 5 post-ruxolitinib treatment. To measure IFN-γ protein, supernatants were diluted at 1:4 and measured by Bio-Plex Pro mouse IFN-γ immunoassay (Bio-Rad) according to manufacturer’s instructions.

### Tumor models

7-10 week-old aged control female C57BL/6 mice were implanted intradermally in the right flank with 1-5x10^5^ cells (MC38 and knockout MC38 cell lines), subcutaneously (s.c.) in the right flank with 5x10^6^ cells (A20), 1x10^6^ (EL4) or 2.5x10^5^ (LLC1). For tumor-infiltrating lymphocyte analysis, mice were treated with 30 mg/kg ruxolitinib phosphate (LC Laboratories cat #R-6688) daily from day 8 until day 15 post-implantation by oral gavage. For TIL isolation, at day 16 post-implantation, tumors were excised, manually dissociated, incubated with collagenase, hyaluronidase and DNase (STEMCELL Technologies cat #07912, 100-0762) according to manufacturer’s instructions and mashed through 100 μm filters. TILs were then stained with surface antibodies and analyzed by flow cytometry. For intranuclear staining, cells were fixed overnight with FOXP3 Fixation/Permeabilization reagent (eBioscience, catalog# 00-5523-00) and stained with nuclear protein targeting antibodies the following day. For blood analysis, blood was isolated using the retroorbital route and red blood cells lysed twice using Lonza ACK Lysing buffer (Lonza, Basel, Switzerland), then subjected to staining as described above. For bone marrow analysis, femur bone marrow was isolated and stained as described above. For splenocyte analysis, spleens were manually dissociated and mashed through 100 μm filters, then stained as described above. Ruxolitinib phosphate treatment was performed using daily oral gavage, concentration 30 mg/kg unless specified otherwise in text. Immune checkpoint inhibitor treatment was performed at the specified timepoint using anti-PD1 (50 μg/mouse, Leinco cat #P362) and anti-CTLA4 (50 μg/mouse, bioXcell cat #BE0164) or isotype controls (mouse IgG2b and rat IgG2a) by intraperitoneal injection. For cell depletion experiments, CD8 T cells, NK cells and granulocytes were depleted using anti-CD8 (Leinco cat #C2442), anti-NK1.1 (bioXcell cat #BE0036) and anti-Ly6G (Leinco cat #L280) antibodies, respectively, or respective control antibodies (rat IgG2b, mouse IgG2a, rat IgG2a) using intraperitoneal injection 1 day before ICI treatment and every 2-3 days thereafter until end of ICI treatment. On-treatment depletion efficiency was verified by flow cytometry of peripheral blood cells.

Cell lines were obtained from ATCC (EL4, A20) or Kerafast (MC38). LLC1 cell line was a kind gift from Dr Linda Sherman. *Jak1*^-/-^ and *Jak2*^-/-^ MC38 cells were kindly provided by Dr Antoni Ribas (*41*) and authenticated by bulk RNA-seq which detected frameshift mutations at the *Jak1* and *Jak2* loci, respectively.

### Myeloid cell functional assays

To test the ability of MDSCs to suppress NK cell proliferation and cytokine production, Gr1^+^ cells from MC38 tumors were isolated using the CD11b^+^ Gr1^+^ magnetic isolation kit (STEMCELL Technologies cat #19867) and incubated with NK cells isolated using the mouse NK cell isolation kit (STEMCELL Technologies cat #19855) from tumor-free wild type B6 mice in the indicated ratios in the presence of IL-2 as described previously (*72*). To test the ability of granulocytes to stimulate T cell proliferation, T cells were isolated from tumor-free wild type B6 mice using the mouse CD8 T cell isolation kit (STEMCELL Technologies cat #19853), labeled with Cell Trace Violet (Thermo Fisher cat #C34557) according to the manufacturer’s specifications; live CD11b^+^ Ly6G^+^ Ly6C^lo^ were isolated from spleens of mice treated with ICI, ICI+rux, ruxolitinib or vehicle and incubated with the indicated ratios of T cells in anti-CD3/CD28 coated plates for 3 days, then analyzed by flow cytometry.

### Immunohistochemistry

Freshly isolated tumor tissue was fixed using Bouin’s solution (Sigma Aldrich) and subjected to tissue processing for paraffin-embedded sectioning accordingly to standard protocols. Immunohistochemistry staining was performed using anti-NK1.1 antibody and 3,3’ Diaminobenzidine (DAB) Substrate Kit.

### High-throughput screen for compounds that rescue T cell exhaustion

An assay using a YFP-IFNγ reporter strain was developed and scaled to high throughput format as described previously (*17*). Briefly, B cell depleted splenocytes were prepared from LCMV-Cl13 infected IFN-γ-YFP mice at day 15 p.i. Next, 5x10^4^ cells were seeded onto 384-well plate wells pre-spotted with DMSO or ReFrame compounds at 5 µM final concentration. Following a 5-day incubation period, the viability dye 7-AAD was added to each well (1:50 dilution) and plates were rested for 15 minutes. Following a 5-day incubation period, the viability dye 7-AAD was added to each well (1:50 dilution) and plates were rested for 15 minutes.

Cell-free IC_50_ values were derived from the sources below.

|  |  |
| --- | --- |
| ruxolitinib | (21) |
| tofacitinib | (73) |
| CHZ868 | (74) |
| AZD-1480 | (75) |
| filgotinib | (76) |
| peficitinib | (77) |
| oclacitinib | (78) |
| TG101209 | (79) |
| NVP-BSK805 | (80) |
| BMS911543 | (81) |

### Flow cytometry

Surface staining: Single cell suspensions were incubated with 5% normal rat serum and rat anti-mouse CD16/CD32 antibody (BD Biosciences close 2.4G2) then stained with fluorophore-conjugated antibodies, washed and analyzed by flow cytometry without fixation. Intracellular staining: spleens from Cl13 infected mice were collected and dissociated manually. Red blood cells were lysed using Lonza ACK Lysing buffer (Lonza, Basel, Switzerland). Splenocytes were stained with LCMV-specific gp-33 tetramer and surface antibodies then fixed overnight with FOXP3 Fixation/Permeabilization reagent (eBioscience, catalog #00-5523-00) and stained with intranuclear antibodies the following day. Stained cells were analyzed using the 5-laser Cytek Aurora (Cytek Biosciences, Fremont, CA).

### Bulk RNA sequencing

For bulk PBMC RNA-seq analysis, RNA was isolated using RNeasy Mini kit (Qiagen), polyA+ library preparation performed and followed by 2x150 bp NovaSeq (Illumina) sequencing. For bulk RNA-seq analysis of sorted cells, RNA was isolated using Arcturus Picopure RNA kit (Thermo), converted to cDNA and subjected to amplification followed by 2x150 bp sequencing on the NovaSeq (Illumina).

### Single-cell RNA-sequencing

Live single cells were isolated from peripheral blood mononuclear cells using flow cytometry. All samples were subjected to 3’ transcriptome single-cell library preparation using Chromium Single Cell Gene Expression kit version 3.1 (10X Genomics, Pleasanton, CA). For CITE-seq experiments, cells were first stained using TotalSeq B DNA barcoded antibodies (Biolegend) including cell hashing antibodies, then subjected to single-cell transcriptome and barcode library preparation using the Chromium Single Cell Gene Expression kit version 3.1 (10X Genomics, Pleasanton, CA). Libraries were sequenced using NextSeq 2000 or NovaSeq (Illumina, San Diego, CA) to an average depth of 10,000 reads per cell.

### Bulk RNA-seq analysis

Reads were aligned to the mouse genome and genic reads quantified using STAR version 2.7.0f (Dobin et al., 2013) and Ensembl version 101 GRCm38 genome and transcriptome annotations. Normalization, differential expression analysis, principal component analysis were performed using R package DESeq2 v1.35.0 (Love et al., 2014); heatmaps were constructed using R package ComplexHeatmap v2.12.0. Gene counts for publicly available studies GSE83978, GSE147910 were obtained from the Sequence Read Archive and analyzed using the same pipeline. Analyses were performed using R v4.2.0 (Team, 2008). Digital cytometry was performed using CIBERSORTx (*82*) in single-cell RNA-seq reference mode. The Seurat large PBMC dataset was used as the reference (*83*).

### Single-cell transcriptome analysis

Raw sequencing data were demultiplexed using cellranger mkfastq v6.1 (10X Genomics). Cellranger count v6.1 (10X Genomics) was used to identify cells, align cDNA reads, identify hashtag barcode – cell barcode reads, count reads per gene and produce a cell-gene count matrix. Filtering of low-quality cells, filtering of doublets, dimensionality reduction, clustering and visualization were performed using Seurat v4.0.1.

## Data availability

Mouse transcriptomic studies will be deposited to GEO. Human transcriptomic studies are available on reasonable request. Sharing of raw human sequencing data is subject to IRB review.

## Acknowledgments

We thank The Scripps Research Institute flow cytometry and genomics cores, The UCSD Moores Cancer Center histology core, University of Minnesota Translational Therapy Laboratory, Tony Mondala and Halley Nguyen for experimental assistance. The authors are also grateful to Sue Kaech for advice on data interpretation, Antoni Ribas for kindly providing MC38 knockout cell lines, Daniel Lazar for sharing primary blood cells and Linda Sherman for sharing LLC1 cells. J. Z. is a recipient of the Cancer Research Institute/Irvington postdoctoral fellowship. This work was funded by National Institute of Health grants R01AI123210, UL1TR002550 Pilot Award, R01AI164744 (to J. R. T.) and R21AI141842 (to L. L.). The clinical trial was funded by Incyte Corporation and Bristol-Myers Squibb, and performed in collaboration with the Big Ten Cancer Research Consortium.

## Author Contributions

J. Z., I. P., B. S. M., M. B. A. O. and J. R. T. designed the project, J. Z., I. P., B. S. M., K. L. M. and R. B. Z. performed experiments, J. Z., I. P., B. S. M. analyzed data, V. B. provided patient samples and data, M. B. A. O., L. L. L. and J. R. T. obtained funding, J. Z. and J. R. T. wrote the paper, all authors interpreted data, all authors reviewed the manuscript.

## Supplementary Figures

**Supplementary Figure 1.**
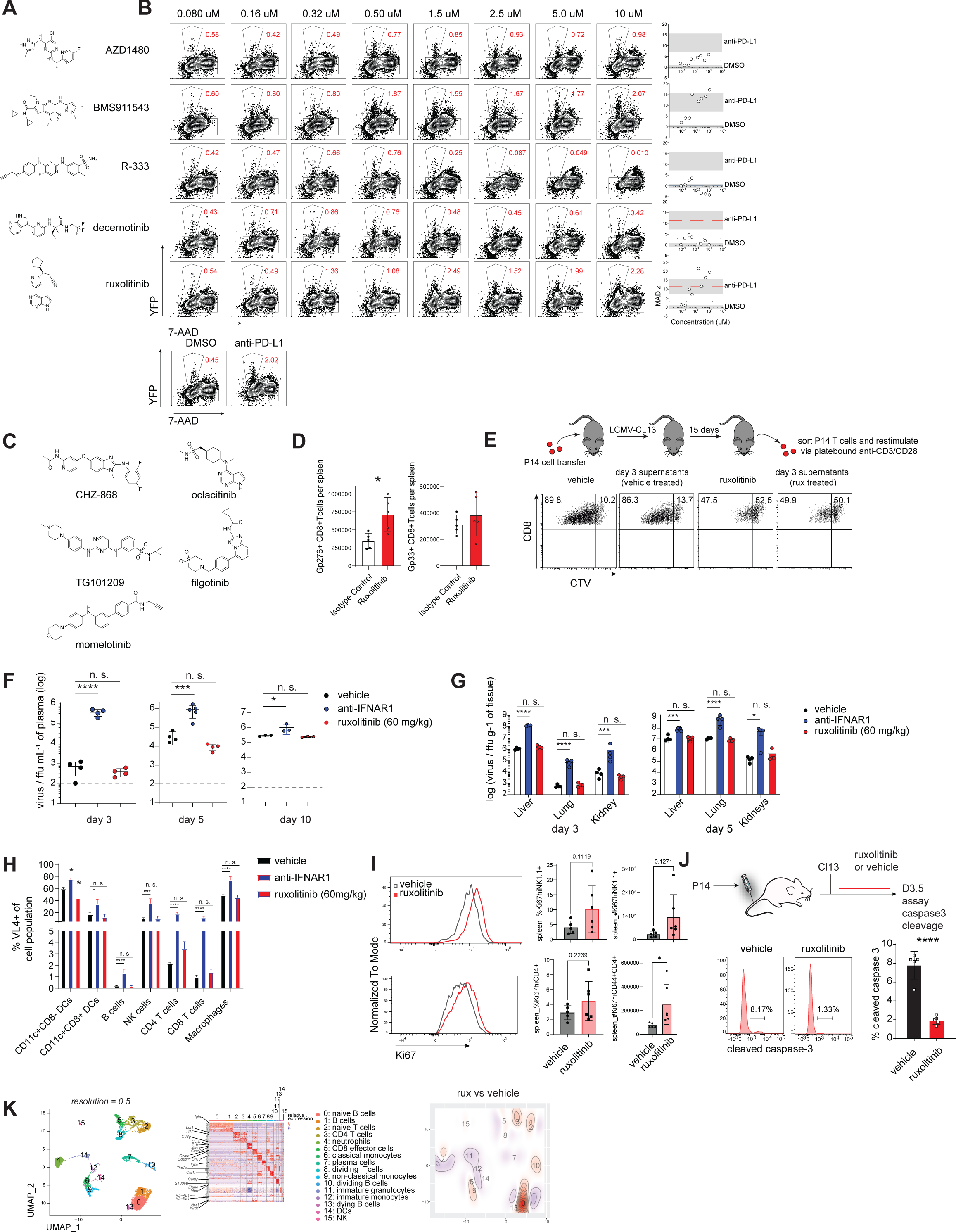
(A) Chemical structures of top JAK inhibitor hits in the ReFrame library screen; (B) results of validation assay of re-obtained JAK inhibitor hits performed as the primary screening assay at a range of concentrations for each compound; (C) chemical structures of top 5 hits from kinase inhibitor library (MedChemExpress); (D) splenic virus-specific CD8 T cells in mice treated with ruxolitinib or vehicle at d8 post infection; (E) P14 cells were adoptively transferred to animals infected with Clone 13, then harvested at 15 dpi, stained with CTV dye and stimulated with plate-bound anti-CD3/CD28 in the presence of ruxolitinib or vehicle for 3 days; (F-G) mice treated with anti-IFNAR1 or isotype 1 day prior to infection with LCMV Cl13 then treated daily with ruxolitinib or vehicle: (G) LCMV Cl13 titres in the organs analyzed at the specified time post infection, (H) percentage of cells in spleen positive for LCMV protein by cell type; (I) splenic lymphocytes at 10 dpi in Cl13 infected mice treated with ruxolitinib or vehicle; (J) cell death in P14 cells adoptively transferred to Cl13 infected animals treated with ruxolitinib or vehicle and analyzed at 3.5 dpi; (K) single-cell transcriptome analysis of splenocytes from Cl13 infected mice treated with ruxolitinib or vehicle, cells analyzed at 10 dpi. Statistical comparison of experimental groups was performed using Student’s two-tailed t-test (pairwise comparisons) or one-way ANOVA with Dunnett’s post-test (multi-group comparisons); bars show standard deviation. *, *p* ≤ 0.05; ****, *p* ≤ 0.0001; n. s., not significant.

**Supplementary Figure 2.**
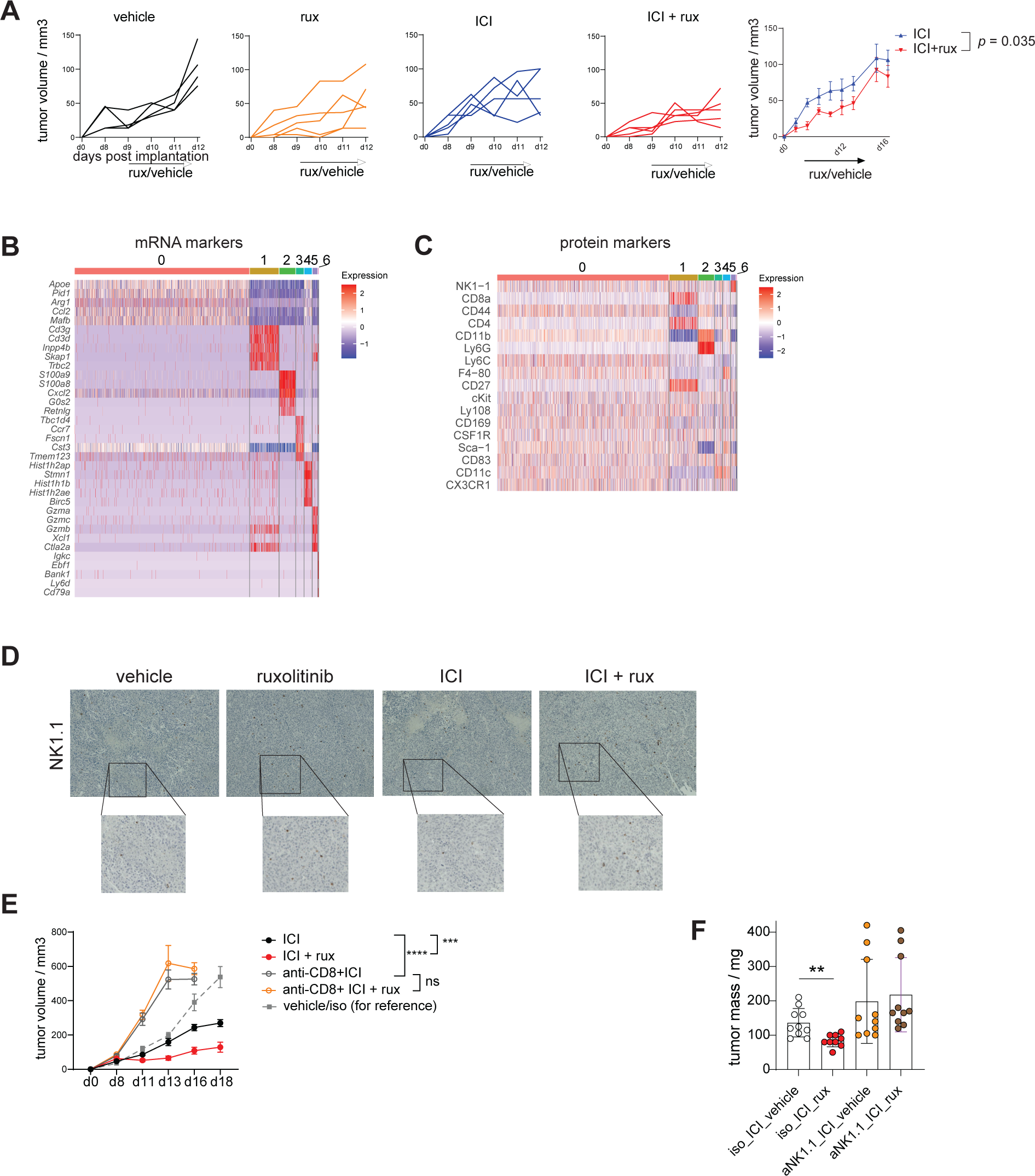
(A) B6 mice received LLC1 tumor cells subcutaneously and were treated with ICI or isotype every 7 days and ruxolitinib or vehicle every day since palpable; (B-C) heatmap of mRNA and protein markers of CITE-seq clusters defined in panel 2C; (D) immunohistochemistry staining of NK1.1 in MC38 tumor sections at d18 post implantation, treatment groups as described in panel 2A; (E) mice bearing MC38 tumors received CD8 T cell depleting antibody or isotype control 2 days prior to treatment with ICI/isotype and ruxolitinib/vehicle as described in Figure 2A, then every 3 days thereafter; (F) tumor weights of mice from NK cell depletion experiment described in Figure 2L. Statistical comparison of experimental groups was performed using two-way ANOVA (A, E) or two-tailed Student’s t-test (F). **, *p* ≤ 0.01; ***, *p* ≤ 0.001; ****, *p* ≤ 0.0001.

**Supplementary Figure 3.**
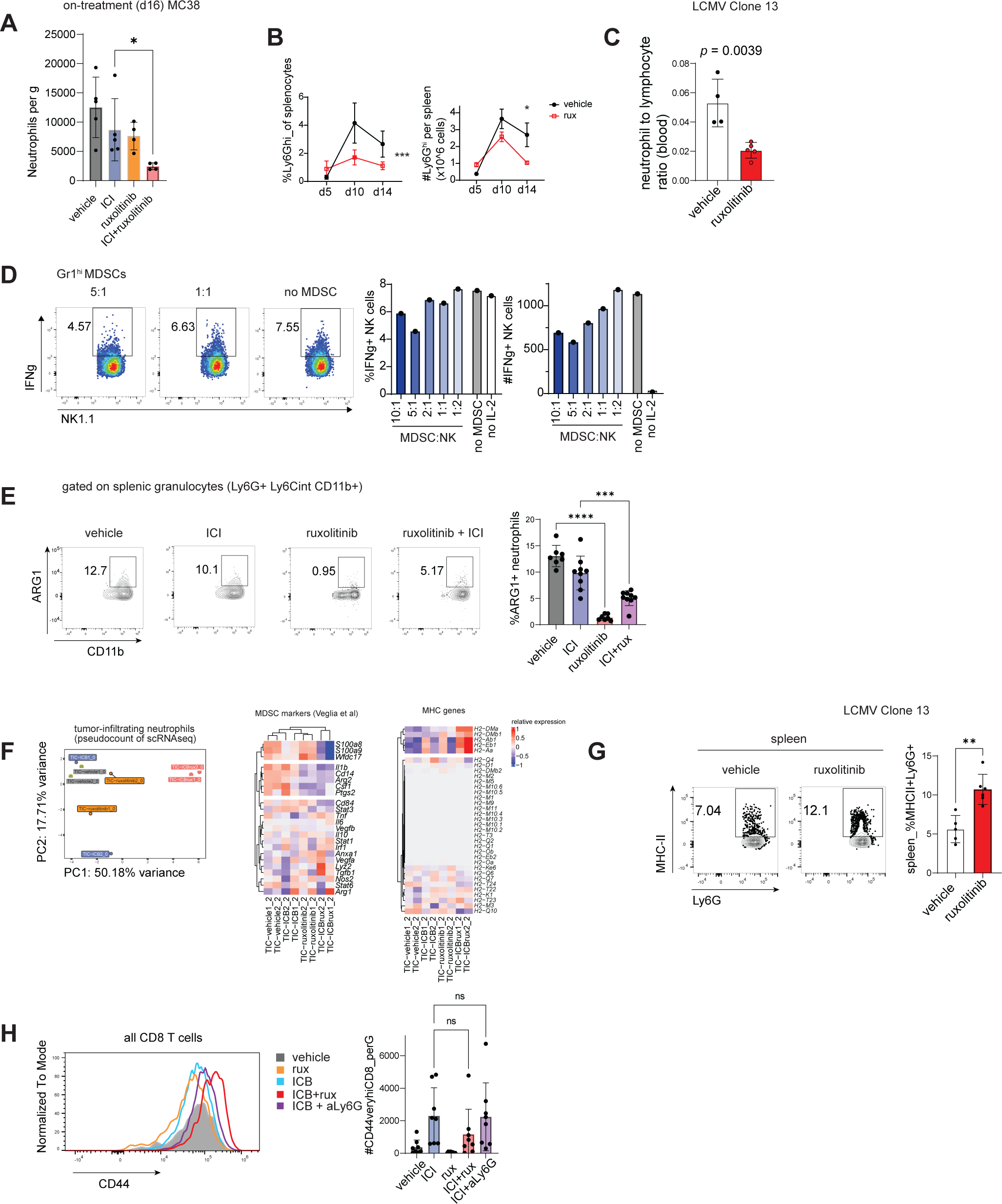
(A) MC38 tumor bearing mice were treated as in Figure 2A, tumor-infiltrating cells analyzed on-treatment at ∼d16 post implantation; (B-C) B6 mice were infected with 2x10^6^ pfu Cl13 and treated with ruxolitinib or vehicle by daily gavage, splenocytes analyzed by flow cytometry at days 5, 10 and 14 (B) or blood analyzed at day 10 (C) post infection; (D) tumor-infiltrating myeloid derived suppressor cells from MC38 tumor bearing mice were isolated and subjected to NK cell suppression assay (*72*); (E) MC38 tumor bearing mice were treated as in Figure 2A, splenic granulocytes analyzed 2 days after treatment cessation; (F) principal component analysis and relative expression of selected genes in RNA-seq dataset described in Figure 3D; (G) splenic granulocytes from Cl13 infected mice treated with ruxolitinib or vehicle at 10 dpi; (H) tumor-infiltrating cells in mice treated as in Figure 3C were analyzed 2 days post treatment. Statistical comparison of experimental groups was performed using one-way ANOVA with Dunnett’s post-test (A, E, H), two-way ANOVA (B) or two-tailed Student’s t-test (C, G); *, *p* ≤ 0.05; **, *p* ≤ 0.01; ***, *p* ≤ 0.001; ****, *p* ≤ 0.0001.

**Supplementary Figure 4.**
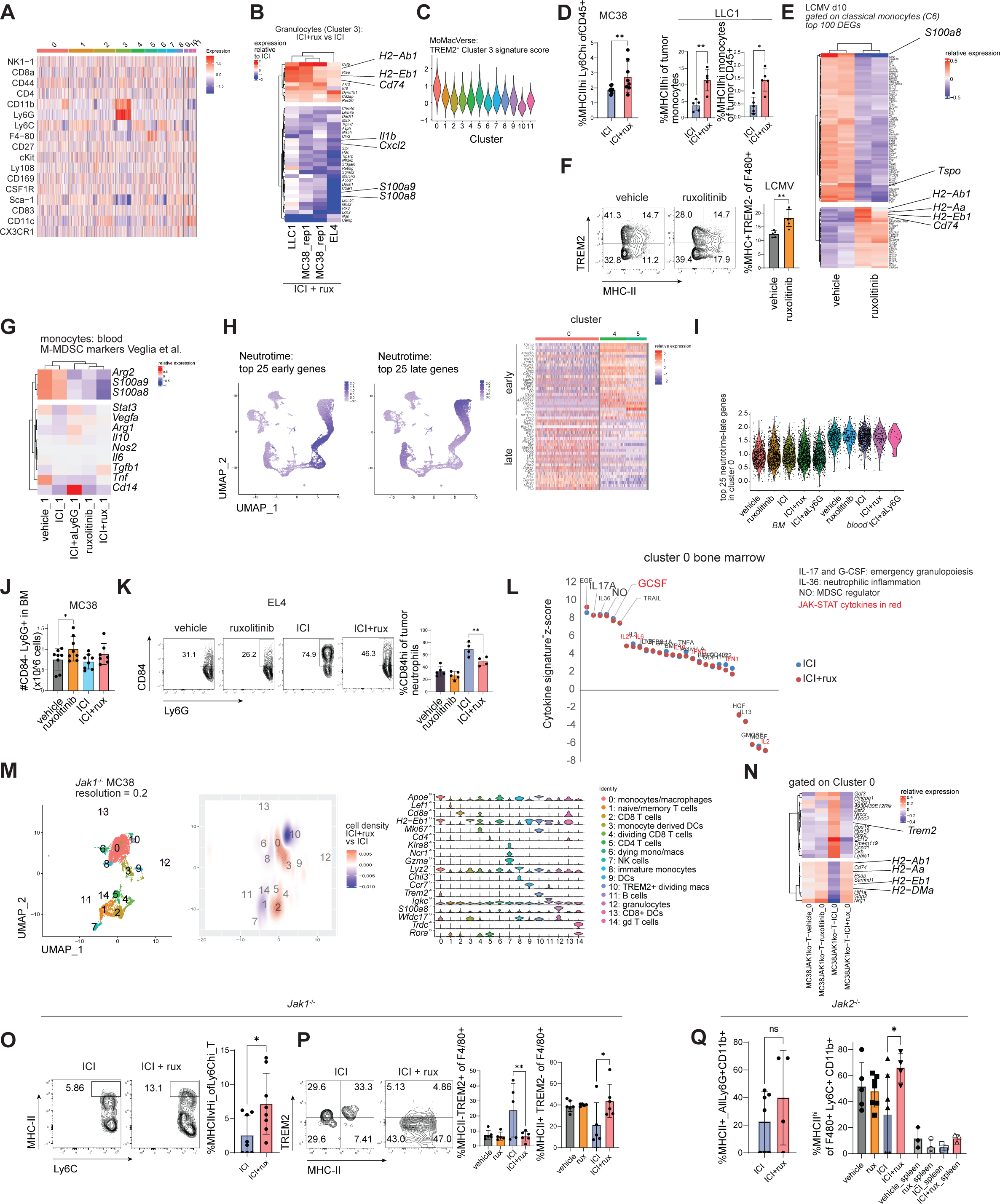
(A) protein markers of CITE-seq dataset in Figure 4A; (B) relative expression of ruxolitinin-regulated genes in tumor-infiltrating granulocytes from ICI+rux treated mice compared to respective ICI treated controls; (C) relative expression of Cluster 3 signature from the MoMac-VERSE among myeloid clusters described in Figure 4A; (D) tumor-infiltrating monocytic cells from mice bearing MC38 or LLC1 tumors and treated as in Figure 2A were analyzed by flow cytometry 2 days after cessation of treatment; (E) top differentially expressed genes in splenic classical monocytes in ruxolitinib vs vehicle treated Clone 13 infected mice 10 dpi, gated on Cluster 6 from Figure S1K; (F) gated on splenic macrophages (F4/80+ Ly6G-CD11b-) in Clone 13 infected mice 10 dpi; (G) relative expression of M-MDSC markers in blood monocytes (Cluster 1 from CITE-seq dataset described in Figure 4G); (H) relative expression of early and late neutrophil differentiation signatures in CITE-seq dataset in Figure 4G, top 25 early and late Neutrotime genes were selected for analysis; (I) relative expression of gene signature comprising top 25 late Neutrotime genes in Cluster 0 of CITE-seq dataset in Figure 4G; (J) total number of CD84^-^ granulocytes in bone marrow of mice treated as in Figure 2A; (K) mice bearing EL4 tumors were treated analogous to Fig. 2A and tumor-infiltrating granulocytes analyzed at size-based endpoint; (L) Cytokine transcriptomic signature scores in Cluster 0 bone marrow cells ranked by CytoSig (*40*), important myeloid and MDSC cytokines highlighted; (M-Q) Mice bearing *Jak1*^-/-^ (M-P) or *Jak2*^-/-^ (Q) MC38 tumors were treated with ICI and ruxolitinib or vehicle analogous to Fig. 2A and tumor-infiltrating granulocytes analyzed 2 days post treatment; (M) single-cell transcriptomic analysis of tumor-infiltrating CD45^+^ cells; (N) genes differentially expressed between ICI+rux and ICI treated groups in Cluster 0; (O) flow cytometry of tumor-infiltrating monocytes (Ly6C^hi^ CD11b^+^ Ly6G^-^); (P) gated on tumor macrophages; (Q) flow cytometry studies in *Jak2*^-/-^ MC38 tumors, gated on granulocytes (left) and macrophages (right). Statistical comparison of experimental groups was performed using Student’s two-tailed t-test (D, F, O, Q left) or one-way ANOVA with Dunnett’s post-test (J, K, P, Q right); *, *p* ≤ 0.05; **, *p* ≤ 0.01; ***, *p* ≤ 0.001; ****, *p* ≤ 0.0001.

**Supplementary Figure 5.**
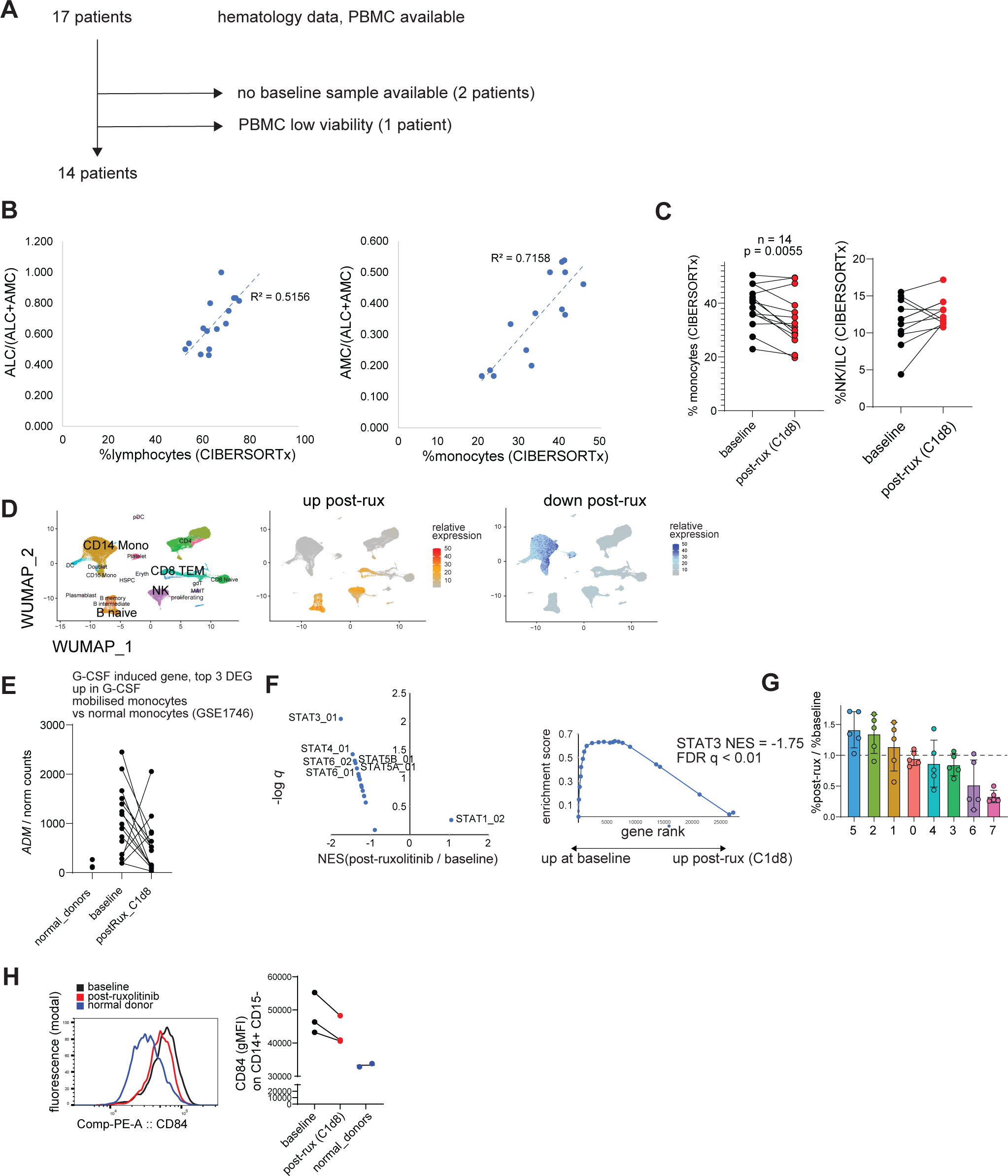
(A) Details on patients included in analysis; (B) peripheral blood cells examined by complete cell count and bulk RNA-sequencing, graph shows comparison of absolute lymphocyte count (ALC) as fraction of absolute monocyte count and ALC combined vs predicted percentage of lymphocyte from RNA-seq (left), AMC as fraction of AMC and ALC combined vs predicted monocyte fraction in PBMC from RNA-seq (right); (C) relative percentage of monocytes (left) and NK/ILCs (right) in PBMC before and after ruxolitinib; (D) expression pattern of ruxolitinib-upregulated and -downregulated genes in a dataset of normal donor PBMCs (*83*); (E) normalized expression of ADM; (F) normalized enrichment scores and q values of STAT gene sets in PBMC bulk-seq data; (G) ruxolitinib-induced changes in individual patient PBMC scRNA-seq datasets; (H) flow cytometric analysis of classical monocytes in patients and normal donors. Statistical comparison of experimental groups was performed using paired Student’s t-test (C).

## References

1. R. Weber, V. Fleming, X. Hu, V. Nagibin, C. Groth, P. Altevogt, J. Utikal, V. Umansky, Myeloid-Derived Suppressor Cells Hinder the Anti-Cancer Activity of Immune Checkpoint Inhibitors. Frontiers in immunology 9, (2018).

2. X. Liu, G. D. Hogg, D. G. DeNardo, Rethinking immune checkpoint blockade: ‘Beyond the T cell’. Journal for ImmunoTherapy of Cancer 9, e001460 (2021).

3. R. S. Hotchkiss, L. L. Moldawer, Parallels between Cancer and Infectious Disease. New England Journal of Medicine 371, 380–383 (2014).

4. E. J. Wherry, M. Kurachi, Molecular and cellular insights into T cell exhaustion. Nature reviews. Immunology 15, 486–499 (2015).

5. D. L. Barber, E. J. Wherry, D. Masopust, B. Zhu, J. P. Allison, A. H. Sharpe, G. J. Freeman, R. Ahmed, Restoring function in exhausted CD8 T cells during chronic viral infection. Nature 439, 682–687 (2006).

6. Brian A. Norris, Luke S. Uebelhoer, Helder I. Nakaya, Aryn A. Price, A. Grakoui, B. Pulendran, Chronic but Not Acute Virus Infection Induces Sustained Expansion of Myeloid Suppressor Cell Numbers that Inhibit Viral-Specific T Cell Immunity. Immunity 38, 309–321 (2013).

7. L. M. Snell, T. L. McGaha, D. G. Brooks, Type I Interferon in Chronic Virus Infection and Cancer. Trends in Immunology 38, 542–557 (2017).

8. S. Hegde, A. M. Leader, M. Merad, MDSC: Markers, development, states, and unaddressed complexity. Immunity 54, 875–884 (2021).

9. F. Veglia, E. Sanseviero, D. I. Gabrilovich, Myeloid-derived suppressor cells in the era of increasing myeloid cell diversity. Nature Reviews Immunology, (2021).

10. N. de Haas, C. de Koning, L. Spilgies, I. J. M. de Vries, S. V. Hato, Improving cancer immunotherapy by targeting the STATe of MDSCs. OncoImmunology 5, e1196312 (2016).

11. A. M. K. Law, F. Valdes-Mora, D. Gallego-Ortega, Myeloid-Derived Suppressor Cells as a Therapeutic Target for Cancer. Cells 9, (2020).

12. S. K. Maenhout, S. Du Four, J. Corthals, B. Neyns, K. Thielemans, J. L. Aerts, AZD1480 delays tumor growth in a melanoma model while enhancing the suppressive activity of myeloid-derived suppressor cells. Oncotarget*; Vol* 5*, No* 16, (2014).

13. S. Sendo, J. Saegusa, H. Yamada, K. Nishimura, A. Morinobu, Tofacitinib facilitates the expansion of myeloid-derived suppressor cells and ameliorates interstitial lung disease in SKG mice. Arthritis Research & Therapy 21, 184 (2019).

14. W. Xiao, J. D. Klement, C. Lu, M. L. Ibrahim, K. Liu, IFNAR1 Controls Autocrine Type I IFN Regulation of PD-L1 Expression in Myeloid-Derived Suppressor Cells. The Journal of Immunology 201, 264 (2018).

15. J. J. O’Shea, D. M. Schwartz, A. V. Villarino, M. Gadina, I. B. McInnes, A. Laurence, The JAK-STAT Pathway: Impact on Human Disease and Therapeutic Intervention. Annual Review of Medicine 66, 311–328 (2015).

16. Y. Tanaka, Y. Luo, J. J. O’Shea, S. Nakayamada, Janus kinase-targeting therapies in rheumatology: a mechanisms-based approach. Nature Reviews Rheumatology 18, 133–145 (2022).

17. B. S. Marro, J. Zak, R. B. Zavareh, J. R. Teijaro, L. L. Lairson, M. B. A. Oldstone, Discovery of Small Molecules for the Reversal of T Cell Exhaustion. Cell Reports 29, 3293–3302.e3293 (2019).

18. K. C. Cotto, A. H. Wagner, Y.-Y. Feng, S. Kiwala, A. C. Coffman, G. Spies, A. Wollam, N. C. Spies, O. L. Griffith, M. Griffith, DGIdb 3.0: a redesign and expansion of the drug–gene interaction database. Nucleic Acids Research 46, D1068–D1073 (2017).

19. C. E. Rudd, K. Chanthong, A. Taylor, Small Molecule Inhibition of GSK-3 Specifically Inhibits the Transcription of Inhibitory Co-receptor LAG-3 for Enhanced Anti-tumor Immunity. Cell Reports 30, 2075–2082.e2074 (2020).

20. Jakafi (ruxolitinib) [package insert]. U. S. Food and Drug Administration website. 2019.

21. A. Quintás-Cardama, K. Vaddi, P. Liu, T. Manshouri, J. Li, P. A. Scherle, E. Caulder, X. Wen, Y. Li, P. Waeltz, M. Rupar, T. Burn, Y. Lo, J. Kelley, M. Covington, S. Shepard, J. D. Rodgers, P. Haley, H. Kantarjian, J. S. Fridman, S. Verstovsek, Preclinical characterization of the selective JAK1/2 inhibitor INCB018424: therapeutic implications for the treatment of myeloproliferative neoplasms. Blood 115, 3109–3117 (2010).

22. F. Lussana, M. Cattaneo, A. Rambaldi, A. Squizzato, Ruxolitinib-associated infections: A systematic review and meta-analysis. American Journal of Hematology 93, 339–347 (2018).

23. K. L. Winthrop, The emerging safety profile of JAK inhibitors in rheumatic disease. Nature Reviews Rheumatology 13, 234–243 (2017).

24. J. R. Teijaro, C. Ng, A. M. Lee, B. M. Sullivan, K. C. F. Sheehan, M. Welch, R. D. Schreiber, J. Carlos de la Torre, M. B. A. Oldstone, Persistent LCMV Infection Is Controlled by Blockade of Type I Interferon Signaling. Science 340, 207 (2013).

25. E. B. Wilson, D. H. Yamada, H. Elsaesser, J. Herskovitz, J. Deng, G. Cheng, B. J. Aronow, C. L. Karp, D. G. Brooks, Blockade of Chronic Type I Interferon Signaling to Control Persistent LCMV Infection. Science 340, 202–207 (2013).

26. E. M. Elli, C. Baratè, F. Mendicino, F. Palandri, G. A. Palumbo, Mechanisms Underlying the Anti-inflammatory and Immunosuppressive Activity of Ruxolitinib. Frontiers in Oncology 9, (2019).

27. E. Perez-Ruiz, L. Minute, I. Otano, M. Alvarez, M. C. Ochoa, V. Belsue, C. de Andrea, M. E. Rodriguez-Ruiz, J. L. Perez-Gracia, I. Marquez-Rodas, C. Llacer, M. Alvarez, V. de Luque, C. Molina, A. Teijeira, P. Berraondo, I. Melero, Prophylactic TNF blockade uncouples efficacy and toxicity in dual CTLA-4 and PD-1 immunotherapy. Nature 569, 428–432 (2019).

28. J. L. Benci, B. Xu, Y. Qiu, T. J. Wu, H. Dada, C. Twyman-Saint Victor, L. Cucolo, D. S. M. Lee, K. E. Pauken, A. C. Huang, T. C. Gangadhar, R. K. Amaravadi, L. M. Schuchter, M. D. Feldman, H. Ishwaran, R. H. Vonderheide, A. Maity, E. J. Wherry, A. J. Minn, Tumor Interferon Signaling Regulates a Multigenic Resistance Program to Immune Checkpoint Blockade. Cell 167, 1540–1554.e1512 (2016).

29. M. Stoeckius, C. Hafemeister, W. Stephenson, B. Houck-Loomis, P. K. Chattopadhyay, H. Swerdlow, R. Satija, P. Smibert, Simultaneous epitope and transcriptome measurement in single cells. Nature Methods 14, 865–868 (2017).

30. V. Bronte, S. Brandau, S.-H. Chen, M. P. Colombo, A. B. Frey, T. F. Greten, S. Mandruzzato, P. J. Murray, A. Ochoa, S. Ostrand-Rosenberg, P. C. Rodriguez, A. Sica, V. Umansky, R. H. Vonderheide, D. I. Gabrilovich, Recommendations for myeloid-derived suppressor cell nomenclature and characterization standards. Nature Communications 7, 12150 (2016).

31. K. Mulder, A. A. Patel, W. T. Kong, C. Piot, E. Halitzki, G. Dunsmore, S. Khalilnezhad, S. E. Irac, A. Dubuisson, M. Chevrier, X. M. Zhang, J. K. C. Tam, T. K. H. Lim, R. M. M. Wong, R. Pai, A. I. S. Khalil, P. K. H. Chow, S. Z. Wu, G. Al-Eryani, D. Roden, A. Swarbrick, J. K. Y. Chan, S. Albani, L. Derosa, L. Zitvogel, A. Sharma, J. Chen, A. Silvin, A. Bertoletti, C. Blériot, C.-A. Dutertre, F. Ginhoux, Cross-tissue single-cell landscape of human monocytes and macrophages in health and disease. Immunity 54, 1883–1900.e1885 (2021).

32. L. Dolcetti, E. Peranzoni, S. Ugel, I. Marigo, A. Fernandez Gomez, C. Mesa, M. Geilich, G. Winkels, E. Traggiai, A. Casati, F. Grassi, V. Bronte, Hierarchy of immunosuppressive strength among myeloid-derived suppressor cell subsets is determined by GM-CSF. European Journal of Immunology 40, 22–35 (2010).

33. S. I. Abrams, J. D. Waight, Identification of a G-CSF-Granulocytic MDSC axis that promotes tumor progression. OncoImmunology 1, 550–551 (2012).

34. R. Grieshaber-Bouyer, F. A. Radtke, P. Cunin, G. Stifano, A. Levescot, B. Vijaykumar, N. Nelson-Maney, R. B. Blaustein, P. A. Monach, P. A. Nigrovic, O. Aguilar, R. Allan, J. Astarita, K. F. Austen, N. Barrett, A. Baysoy, C. Benoist, B. D. Brown, M. Buechler, J. Buenrostro, M. A. Casanova, K. Chowdhary, M. Colonna, T. Crowl, T. Deng, F. Desland, M. Dhainaut, J. Ding, C. Dominguez, D. Dwyer, M. Frascoli, S. Gal-Oz, A. Goldrath, T. Johanson, S. Jordan, J. Kang, V. Kapoor, E. Kenigsberg, J. Kim, K. w. Kim, E. Kiner, M. Kronenberg, L. Lanier, C. Laplace, C. Lareau, A. Leader, J. Lee, A. Magen, B. Maier, A. Maslova, D. Mathis, A. McFarland, M. Merad, E. Meunier, P. A. Monach, S. Mostafavi, S. Muller, C. Muus, H. Ner-Gaon, Q. Nguyen, G. Novakovsky, S. Nutt, K. Omilusik, A. Ortiz-Lopez, M. Paynich, V. Peng, M. Potempa, R. Pradhan, S. Quon, R. Ramirez, D. Ramanan, G. Randolph, A. Regev, S. A. Rose, K. Seddu, T. Shay, A. Shemesh, J. Shyer, C. Smilie, N. Spidale, A. Subramanian, K. Sylvia, J. Tellier, S. Turley, B. Vijaykumar, A. Wagers, C. Wang, P. L. Wang, A. Wroblewska, L. Yang, A. Yim, H. Yoshida, C. ImmGen, The neutrotime transcriptional signature defines a single continuum of neutrophils across biological compartments. Nature Communications 12, 2856 (2021).

35. H. Alshetaiwi, N. Pervolarakis, L. McIntyre Laura, D. Ma, Q. Nguyen, A. Rath Jan, K. Nee, G. Hernandez, K. Evans, L. Torosian, A. Silva, C. Walsh, K. Kessenbrock, Defining the emergence of myeloid-derived suppressor cells in breast cancer using single-cell transcriptomics. Science Immunology 5, eaay6017 (2020).

36. B. Yu, L. Yang, S. Song, W. Li, H. Wang, J. Cheng, LRG1 facilitates corneal fibrotic response by inducing neutrophil chemotaxis via Stat3 signaling in alkali-burned mouse corneas. American Journal of Physiology-Cell Physiology 321, C415–C428 (2021).

37. C. B. de Lima, E. K. Tamura, T. Montero-Melendez, J. Palermo-Neto, M. Perretti, R. P. Markus, S. H. P. Farsky, Actions of translocator protein ligands on neutrophil adhesion and motility induced by G-protein coupled receptor signaling. Biochemical and Biophysical Research Communications 417, 918–923 (2012).

38. F. Shojaei, X. Wu, C. Zhong, L. Yu, X.-H. Liang, J. Yao, D. Blanchard, C. Bais, F. V. Peale, N. van Bruggen, C. Ho, J. Ross, M. Tan, R. A. D. Carano, Y. G. Meng, N. Ferrara, Bv8 regulates myeloid-cell-dependent tumour angiogenesis. Nature 450, 825–831 (2007).

39. V. F. Curtis, H. Wang, P. Yang, R. E. McLendon, X. Li, Q.-Y. Zhou, X.-F. Wang, A PK2/Bv8/PROK2 Antagonist Suppresses Tumorigenic Processes by Inhibiting Angiogenesis in Glioma and Blocking Myeloid Cell Infiltration in Pancreatic Cancer. PLOS ONE 8, e54916 (2013).

40. P. Jiang, Y. Zhang, B. Ru, Y. Yang, T. Vu, R. Paul, A. Mirza, G. Altan-Bonnet, L. Liu, E. Ruppin, L. Wakefield, K. W. Wucherpfennig, Systematic investigation of cytokine signaling activity at the tissue and single-cell levels. Nature Methods 18, 1181–1191 (2021).

41. D. Y. Torrejon, G. Abril-Rodriguez, A. S. Champhekar, J. Tsoi, K. M. Campbell, A. Kalbasi, G. Parisi, J. M. Zaretsky, A. Garcia-Diaz, C. Puig-Saus, G. Cheung-Lau, T. Wohlwender, P. Krystofinski, A. Vega-Crespo, C. M. Lee, P. Mascaro, C. S. Grasso, B. Berent-Maoz, B. Comin-Anduix, S. Hu-Lieskovan, A. Ribas, Overcoming Genetically Based Resistance Mechanisms to PD-1 Blockade. Cancer Discovery 10, 1140–1157 (2020).

42. F. Z. Cader, X. Hu, W. L. Goh, K. Wienand, J. Ouyang, E. Mandato, R. Redd, L. N. Lawton, P.-H. Chen, J. L. Weirather, R. C. J. Schackmann, B. Li, W. Ma, P. Armand, S. J. Rodig, D. Neuberg, X. S. Liu, M. A. Shipp, A peripheral immune signature of responsiveness to PD-1 blockade in patients with classical Hodgkin lymphoma. Nature Medicine 26, 1468–1479 (2020).

43. R. Howard, P. A. Kanetsky, K. M. Egan, Exploring the prognostic value of the neutrophil-to-lymphocyte ratio in cancer. Scientific Reports 9, 19673 (2019).

44. A. J. Templeton, M. G. McNamara, B. Šeruga, F. E. Vera-Badillo, P. Aneja, A. Ocaña, R. Leibowitz-Amit, G. Sonpavde, J. J. Knox, B. Tran, I. F. Tannock, E. Amir, Prognostic Role of Neutrophil-to-Lymphocyte Ratio in Solid Tumors: A Systematic Review and Meta-Analysis. JNCI: Journal of the National Cancer Institute 106, (2014).

45. A. Romano, N. L. Parrinello, C. Vetro, A. Chiarenza, C. Cerchione, M. Ippolito, G. A. Palumbo, F. Di Raimondo, Prognostic meaning of neutrophil to lymphocyte ratio (NLR) and lymphocyte to monocyte ration (LMR) in newly diagnosed Hodgkin lymphoma patients treated upfront with a PET-2 based strategy. Annals of Hematology 97, 1009–1018 (2018).

46. V. Bachanova, L. Hegerova, Q. Cao, M. Janakiram, J. Maakaron, S. Ayyappan, D. J. Weisdorf, J. Zak, U. Farooq, V. P. Kenkre, Ruxolitinib Plus Nivolumab in Patients with R/R Hodgkin Lymphoma after Failure of Check-Point Inhibitors: Preliminary Report on Safety and Efficacy. Blood 138, 230 (2021).

47. N. Eric Van Den, A. Marc, G. Thomas, S. Aspasia, H. Corinne, B. Amine, R. Oumedaly, C. Olivier, G. Hervé, V. Gregor, C. Marie-José, A. P. Hélène, C. Marie-Christine, D. Romain, V. Peter, S. Ioanna-Andrea, S. C. Anne, B. Sarah, K. Laurent, M. Franck, A phase II study of the oral JAK1/JAK2 inhibitor ruxolitinib in advanced relapsed/refractory Hodgkin lymphoma. Haematologica 103, 840–848 (2018).

48. S. J. Kim, D. H. Yoon, H. J. Kang, J. Y. Hong, H. S. Lee, S. Y. Oh, H.-J. Shin, J. H. Kong, J. H. Yi, K. Sakamoto, Y. H. Ko, J. Huh, S.-S. Lee, K. Takeuchi, D.-Y. Shin, C. Suh, W. S. Kim, Ruxolitinib shows activity against Hodgkin lymphoma but not primary mediastinal large B-cell lymphoma. BMC Cancer 19, 1080 (2019).

49. P. Armand, A. Engert, A. Younes, M. Fanale, A. Santoro, P. L. Zinzani, J. M. Timmerman, G. P. Collins, R. Ramchandren, J. B. Cohen, J. P. De Boer, J. Kuruvilla, K. J. Savage, M. Trneny, M. A. Shipp, K. Kato, A. Sumbul, B. Farsaci, S. M. Ansell, Nivolumab for Relapsed/Refractory Classic Hodgkin Lymphoma After Failure of Autologous Hematopoietic Cell Transplantation: Extended Follow-Up of the Multicohort Single-Arm Phase II CheckMate 205 Trial. Journal of Clinical Oncology 36, 1428–1439 (2018).

50. A. Heine, S. A. E. Held, S. N. Daecke, S. Wallner, S. P. Yajnanarayana, C. Kurts, D. Wolf, P. Brossart, The JAK-inhibitor ruxolitinib impairs dendritic cell function in vitro and in vivo. Blood 122, 1192 (2013).

51. K. Schönberg, J. Rudolph, M. Vonnahme, S. Parampalli Yajnanarayana, I. Cornez, M. Hejazi, A. R. Manser, M. Uhrberg, W. Verbeek, S. Koschmieder, T. H. Brümmendorf, P. Brossart, A. Heine, D. Wolf, JAK Inhibition Impairs NK Cell Function in Myeloproliferative Neoplasms. Cancer Research 75, 2187–2199 (2015).

52. S. Parampalli Yajnanarayana, T. Stübig, I. Cornez, H. Alchalby, K. Schönberg, J. Rudolph, I. Triviai, C. Wolschke, A. Heine, P. Brossart, N. Kröger, D. Wolf, JAK1/2 inhibition impairs T cell function in vitro and in patients with myeloproliferative neoplasms. British Journal of Haematology 169, 824–833 (2015).

53. D. S. Shin, J. M. Zaretsky, H. Escuin-Ordinas, A. Garcia-Diaz, S. Hu-Lieskovan, A. Kalbasi, C. S. Grasso, W. Hugo, S. Sandoval, D. Y. Torrejon, N. Palaskas, G. A. Rodriguez, G. Parisi, A. Azhdam, B. Chmielowski, G. Cherry, E. Seja, B. Berent-Maoz, I. P. Shintaku, D. T. Le, D. M. Pardoll, L. A. Diaz, Jr., P. C. Tumeh, T. G. Graeber, R. S. Lo, B. Comin-Anduix, A. Ribas, Primary Resistance to PD-1 Blockade Mediated by JAK1/2 Mutations. Cancer Discovery 7, 188–201 (2017).

54. B. Pan, L. Shang, C. Liu, J. Gao, F. Zhang, M. Xu, L. Li, Z. Sun, Z. Li, K. Xu, PD-1 antibody and ruxolitinib enhances graft-versus-lymphoma effect without increasing acute graft-versus-host disease in mice. American Journal of Transplantation 21, 503–514 (2021).

55. C. Lu, A. Talukder, N. M. Savage, N. Singh, K. Liu, JAK-STAT-mediated chronic inflammation impairs cytotoxic T lymphocyte activation to decrease anti-PD-1 immunotherapy efficacy in pancreatic cancer. OncoImmunology 6, e1291106 (2017).

56. R. Ramchandren, E. Domingo-Domènech, A. Rueda, M. Trněný, T. A. Feldman, H. J. Lee, M. Provencio, C. Sillaber, J. B. Cohen, K. J. Savage, W. Willenbacher, A. H. Ligon, J. Ouyang, R. Redd, S. J. Rodig, M. A. Shipp, M. Sacchi, A. Sumbul, P. Armand, S. M. Ansell, Nivolumab for Newly Diagnosed Advanced-Stage Classic Hodgkin Lymphoma: Safety and Efficacy in the Phase II CheckMate 205 Study. Journal of Clinical Oncology 37, 1997–2007 (2019).

57. M. R. Green, S. Monti, S. J. Rodig, P. Juszczynski, T. Currie, E. O’Donnell, B. Chapuy, K. Takeyama, D. Neuberg, T. R. Golub, J. L. Kutok, M. A. Shipp, Integrative analysis reveals selective 9p24.1 amplification, increased PD-1 ligand expression, and further induction via JAK2 in nodular sclerosing Hodgkin lymphoma and primary mediastinal large B-cell lymphoma. Blood 116, 3268–3277 (2010).

58. M. G. M. Roemer, R. H. Advani, A. H. Ligon, Y. Natkunam, R. A. Redd, H. Homer, C. F. Connelly, H. H. Sun, S. E. Daadi, G. J. Freeman, P. Armand, B. Chapuy, D. de Jong, R. T. Hoppe, D. S. Neuberg, S. J. Rodig, M. A. Shipp, PD-L1 and PD-L2 Genetic Alterations Define Classical Hodgkin Lymphoma and Predict Outcome. Journal of Clinical Oncology 34, 2690–2697 (2016).

59. P. E. Debureaux, J. Arrondeau, D. Bouscary, F. Goldwasser, Nivolumab combined with ruxolitinib: antagonism or synergy? Annals of Oncology 29, 1334–1335 (2018).

60. M. Nijland, T. van Meerten, A. Seitz, G. Huls, R. Kibbelaar, L. Visser, A. van den Berg, A. Diepstra, Combined PD-1 and JAK1/2 inhibition in refractory primary mediastinal B-cell lymphoma. Annals of Hematology 97, 905–907 (2018).

61. S. Albeituni, K. C. Verbist, P. E. Tedrick, H. Tillman, J. Picarsic, R. Bassett, K. E. Nichols, Mechanisms of action of ruxolitinib in murine models of hemophagocytic lymphohistiocytosis. Blood 134, 147–159 (2019).

62. V. Kumar, S. Patel, E. Tcyganov, D. I. Gabrilovich, The Nature of Myeloid-Derived Suppressor Cells in the Tumor Microenvironment. Trends in Immunology 37, 208–220 (2016).

63. A. Romano, C. Pavoni, F. Di Raimondo, C. Tarella, S. Viviani, A. Rossi, C. Patti, M. Picardi, M. Cantonetti, G. La Nasa, L. Trentin, S. Bolis, V. Zoli, P. Gavarotti, P. Corradini, M. Cimminiello, C. Schiavotto, G. Parvis, R. Zanotti, G. Gini, A. J. M. Ferreri, P. Viero, S. Chauvie, A. Biggi, A. Massimo Gianni, A. Gallamini, A. Rambaldi, The neutrophil to lymphocyte ratio (NLR) and the presence of large nodal mass are independent predictors of early response: A subanalysis of the prospective phase II PET-2-adapted HD0607 trial. Cancer Medicine 9, 8735–8746 (2020).

64. D. B. Sacdalan, J. A. Lucero, D. L. Sacdalan, Prognostic utility of baseline neutrophil-to-lymphocyte ratio in patients receiving immune checkpoint inhibitors: a review and meta-analysis. Onco Targets Ther 11, 955–965 (2018).

65. M. Hwang, J. V. Canzoniero, S. Rosner, G. Zhang, J. R. White, Z. Belcaid, C. Cherry, A. Balan, G. Pereira, A. Curry, N. Niknafs, J. Zhang, K. N. Smith, L. Sivapalan, J. E. Chaft, J. E. Reuss, K. Marrone, J. C. Murray, Q. K. Li, V. Lam, B. P. Levy, C. Hann, V. E. Velculescu, J. R. Brahmer, P. M. Forde, T. Seiwert, V. Anagnostou, Peripheral blood immune cell dynamics reflect antitumor immune responses and predict clinical response to immunotherapy. Journal for ImmunoTherapy of Cancer 10, e004688 (2022).

66. H. Pircher, K. Burki, R. Lang, H. Hengartner, R. M. Zinkernagel, Tolerance induction in double specific T-cell receptor transgenic mice varies with antigen. Nature 342, 559–561 (1989).

67. R. M. Welsh, M. O. Seedhom, Lymphocytic Choriomeningitis Virus (LCMV): Propagation, Quantitation, and Storage. Current Protocols in Microbiology 8, 15A.11.11–15A.11.11 (2008).

68. M. Battegay, S. Cooper, A. Althage, J. Banziger, H. Hengartner, R. M. Zinkernagel, Quantification of lymphocytic choriomeningitis virus with an immunological focus assay in 24- or 96-well plates. J. Virol. Methods 33, 191–198 (1991).

69. J. Janes, M. E. Young, E. Chen, N. H. Rogers, S. Burgstaller-Muehlbacher, L. D. Hughes, M. S. Love, M. V. Hull, K. L. Kuhen, A. K. Woods, S. B. Joseph, H. M. Petrassi, C. W. McNamara, M. S. Tremblay, A. I. Su, P. G. Schultz, A. K. Chatterjee, The ReFRAME library as a comprehensive drug repurposing library and its application to the treatment of cryptosporidiosis. Proceedings of the National Academy of Sciences 115, 10750 (2018).

70. J. E. Gairin, H. Mazarguil, D. Hudrisier, M. B. Oldstone, Optimal lymphocytic choriomeningitis virus sequences restricted by H-2Db major histocompatibility complex class I molecules and presented to cytotoxic T lymphocytes. Journal of Virology 69, 2297 (1995).

71. A. Oxenius, M. F. Bachmann, P. G. Ashton-Rickardt, S. Tonegawa, R. M. Zinkernagel, H. Hengartner, Presentation of endogenous viral proteins in association with major histocompatibility complex class II: On the role of intracellular compartmentalization, invariant chain and the TAP transporter system. European Journal of Immunology 25, 3402–3411 (1995).

72. S. Greene, Y. Robbins, W. K. Mydlarz, A. P. Huynh, N. C. Schmitt, J. Friedman, L. A. Horn, C. Palena, J. Schlom, D. Y. Maeda, J. A. Zebala, P. E. Clavijo, C. Allen, Inhibition of MDSC Trafficking with SX-682, a CXCR1/2 Inhibitor, Enhances NK-Cell Immunotherapy in Head and Neck Cancer Models. Clinical Cancer Research 26, 1420 (2020).

73. P. S. Changelian, M. E. Flanagan, D. J. Ball, C. R. Kent, K. S. Magnuson, W. H. Martin, B. J. Rizzuti, P. S. Sawyer, B. D. Perry, W. H. Brissette, S. P. McCurdy, E. M. Kudlacz, M. J. Conklyn, E. A. Elliott, E. R. Koslov, M. B. Fisher, T. J. Strelevitz, K. Yoon, D. A. Whipple, J. Sun, M. J. Munchhof, J. L. Doty, J. M. Casavant, T. A. Blumenkopf, M. Hines, M. F. Brown, B. M. Lillie, C. Subramanyam, C. Shang-Poa, A. J. Milici, G. E. Beckius, J. D. Moyer, C. Su, T. G. Woodworth, A. S. Gaweco, C. R. Beals, B. H. Littman, D. A. Fisher, J. F. Smith, P. Zagouras, H. A. Magna, M. J. Saltarelli, K. S. Johnson, L. F. Nelms, S. G. Des Etages, L. S. Hayes, T. T. Kawabata, D. Finco-Kent, D. L. Baker, M. Larson, M.-S. Si, R. Paniagua, J. Higgins, B. Holm, B. Reitz, Y.-J. Zhou, R. E. Morris, J. J. Shea, D. C. Borie, Prevention of Organ Allograft Rejection by a Specific Janus Kinase 3 Inhibitor. Science 302, 875 (2003).

74. S.-C. Wu, Loretta S. Li, N. Kopp, J. Montero, B. Chapuy, A. Yoda, Amanda L. Christie, H. Liu, A. Christodoulou, D. van Bodegom, J. van der Zwet, Jacob V. Layer, T. Tivey, Andrew A. Lane, Jeremy A. Ryan, Samuel Y. Ng, Daniel J. DeAngelo, Richard M. Stone, D. Steensma, M. Wadleigh, M. Harris, E. Mandon, N. Ebel, R. Andraos, V. Romanet, A. Dölemeyer, D. Sterker, M. Zender, Scott J. Rodig, M. Murakami, F. Hofmann, F. Kuo, Michael J. Eck, Lewis B. Silverman, Stephen E. Sallan, A. Letai, F. Baffert, E. Vangrevelinghe, T. Radimerski, C. Gaul, David M. Weinstock, Activity of the Type II JAK2 Inhibitor CHZ868 in B Cell Acute Lymphoblastic Leukemia. Cancer Cell 28, 29–41 (2015).

75. M. Hedvat, D. Huszar, A. Herrmann, J. M. Gozgit, A. Schroeder, A. Sheehy, R. Buettner, D. Proia, C. M. Kowolik, H. Xin, B. Armstrong, G. Bebernitz, S. Weng, L. Wang, M. Ye, K. McEachern, H. Chen, D. Morosini, K. Bell, M. Alimzhanov, S. Ioannidis, P. McCoon, Z. A. Cao, H. Yu, R. Jove, M. Zinda, The JAK2 Inhibitor AZD1480 Potently Blocks Stat3 Signaling and Oncogenesis in Solid Tumors. Cancer Cell 16, 487–497 (2009).

76. L. Van Rompaey, R. Galien, E. M. van der Aar, P. Clement-Lacroix, L. Nelles, B. Smets, L. Lepescheux, T. Christophe, K. Conrath, N. Vandeghinste, B. Vayssiere, S. De Vos, S. Fletcher, R. Brys, G. van ’t Klooster, J. H. M. Feyen, C. Menet, Preclinical Characterization of GLPG0634, a Selective Inhibitor of JAK1, for the Treatment of Inflammatory Diseases. The Journal of Immunology 191, 3568 (2013).

77. M. Ito, S. Yamazaki, K. Yamagami, M. Kuno, Y. Morita, K. Okuma, K. Nakamura, N. Chida, M. Inami, T. Inoue, S. Shirakami, Y. Higashi, A novel JAK inhibitor, peficitinib, demonstrates potent efficacy in a rat adjuvant-induced arthritis model. Journal of Pharmacological Sciences 133, 25–33 (2017).

78. A. J. Gonzales, J. W. Bowman, G. J. Fici, M. Zhang, D. W. Mann, M. Mitton-Fry, Oclacitinib (APOQUEL®) is a novel Janus kinase inhibitor with activity against cytokines involved in allergy. Journal of Veterinary Pharmacology and Therapeutics 37, 317–324 (2014).

79. A. Pardanani, J. Hood, T. Lasho, R. L. Levine, M. B. Martin, G. Noronha, C. Finke, C. C. Mak, R. Mesa, H. Zhu, R. Soll, D. G. Gilliland, A. Tefferi, TG101209, a small molecule JAK2-selective kinase inhibitor potently inhibits myeloproliferative disorder-associated JAK2V617F and MPLW515L/K mutations. Leukemia 21, 1658–1668 (2007).

80. F. Baffert, C. H. Régnier, A. De Pover, C. Pissot-Soldermann, G. A. Tavares, F. Blasco, J. Brueggen, P. Chène, P. Drueckes, D. Erdmann, P. Furet, M. Gerspacher, M. Lang, D. Ledieu, L. Nolan, S. Ruetz, J. Trappe, E. Vangrevelinghe, M. Wartmann, L. Wyder, F. Hofmann, T. Radimerski, Potent and Selective Inhibition of Polycythemia by the Quinoxaline JAK2 Inhibitor NVP-BSK805. Molecular Cancer Therapeutics 9, 1945 (2010).

81. A. Pardanani, A. W. Roberts, J. F. Seymour, K. Burbury, S. Verstovsek, H. M. Kantarjian, K. Begna, H. Yoshitsugu, T. A. Gestone, P. Phillips, G. Xing, G. Peltz, M. V. Lorenzi, L. Alland, A. Woolfson, A. Tefferi, BMS-911543, A Selective JAK2 Inhibitor: A Multicenter Phase 1/2a Study In Myelofibrosis. Blood 122, 664 (2013).

82. A. M. Newman, C. B. Steen, C. L. Liu, A. J. Gentles, A. A. Chaudhuri, F. Scherer, M. S. Khodadoust, M. S. Esfahani, B. A. Luca, D. Steiner, M. Diehn, A. A. Alizadeh, Determining cell type abundance and expression from bulk tissues with digital cytometry. Nature Biotechnology 37, 773–782 (2019).

83. Y. Hao, S. Hao, E. Andersen-Nissen, W. M. Mauck, S. Zheng, A. Butler, M. J. Lee, A. J. Wilk, C. Darby, M. Zager, P. Hoffman, M. Stoeckius, E. Papalexi, E. P. Mimitou, J. Jain, A. Srivastava, T. Stuart, L. M. Fleming, B. Yeung, A. J. Rogers, J. M. McElrath, C. A. Blish, R. Gottardo, P. Smibert, R. Satija, Integrated analysis of multimodal single-cell data. Cell, (2021).

84. A. Subramanian, P. Tamayo, V. K. Mootha, S. Mukherjee, B. L. Ebert, M. A. Gillette, A. Paulovich, S. L. Pomeroy, T. R. Golub, E. S. Lander, J. P. Mesirov, Gene set enrichment analysis: A knowledge-based approach for interpreting genome-wide expression profiles. Proc. Natl Acad. Sci. 102, 15545–15550 (2005).

